# Region Selective Cortical Control Of The Thalamic Reticular Nucleus

**DOI:** 10.1101/2022.01.17.476335

**Authors:** Nóra Hádinger, Emília Bősz, Boglárka Tóth, Gil Vantomme, Anita Lüthi, László Acsády

**Author notes:** Corresponding authors László Acsády Nóra Hádinger.

## Abstract

Corticothalamic pathways, responsible for the top-down control of the thalamus display a classical, canonical organization in that every cortical region sends dual, layer 6 (L6) and layer 5 (L5) output to the thalamus. Here we demonstrate a qualitative, region-specific difference in the organization of corticothalamic pathways. We show that L5 pyramidal cells of the frontal, but not other cortical regions establish monosynaptic connection with the inhibitory thalamic reticular nucleus (TRN). The frontal L5-TRN pathway paralleled the L6-TRN projection but displayed distinct morphological and physiological features. The exact spike output of the L5 contacted TRN cells correlated with the level of cortical synchrony. Optogenetic perturbation of the L5-TRN connection disrupted the tight link between cortical and TRN activity. L5-driven TRN cells innervated all thalamic nuclei involved in the control of frontal cortical activity. Our data show that frontal cortical functions require a highly specialized cortical control over intrathalamic inhibitory processes.

## Introduction

Thalamocortical circuits underlie the organization of all complex behavior. Every cortical region forms tightly organized, bidirectional connections with the thalamus (Deschenes et al., 1998; Guillery and Sherman, 2002; Jones, 2007; Usrey and Sherman, 2019) thus the thalamus forms an integral part of the cortical network (Guo et al., 2017). Thalamocortical circuits involve the excitatory corticothalamic and thalamocortical cells (Jones, 2007) as well as the GABAergic thalamic reticular nucleus (TRN), which is the main source of the intrathalamic inhibition (Halassa and Acsády, 2016; Pinault, 2004). Corticothalamic circuits are regarded as canonical elements of the forebrain in that no qualitative differences are known to be present between different cortical regions (Harris and Shepherd, 2015). In contrast to this view in this study we identified and characterized a region-specific cortico-TRN pathway which arises specifically from the L5 of the frontal cortices.

TRN was shown to be involved in various behavioral processes as sensation (Lee et al., 1994; Le Masson et al., 2002; McCormick and Bal, 1994; Soto-Sánchez et al., 2017), arousal and sleep (Barthó et al., 2014; Herrera et al., 2016; Lewis et al., 2015; Liu et al., 2021; Steriade et al., 1997; Vantomme et al., 2019), selective attention (Ahrens et al., 2015; Crabtree, 2018; Wimmer et al., 2015), spatial navigation (Vantomme et al., 2020), sensory induced flight responses (Dong et al., 2019) and extinction of cued fear conditioning (Lee et al., 2019). It is also involved in a wide range of pathologies like attention-deficit hyperactivity disorder, autism (Wells et al., 2016), epilepsy (Paz et al., 2010; Steriade, 2005) and schizophrenia (Ferrarelli and Tononi, 2011; Steullet et al., 2018). Therefore, understanding the regulation of TRN activity and mapping its possible region- and behavior-specific aspects are crucial to clarify the basics of thalamocortical functions.

The TRN is at the crossroad of thalamocortical circuits. It receives dense topographic input from the thalamocortical cells and is also contacted by excitatory inputs from corticothalamic cells of all cortical areas (Jones, 1975). The top-down cortical inputs to TRN are formed by the collaterals of the layer 6 (L6) corticothalamic cells (Kakei et al., 2001; Pinault et al., 1995). Consequently, TRN can control the direct effect of the L6 activity on the thalamus in a feedforward inhibitory manner (Paz et al., 2011).

The second corticothalamic pathway involves layer 5 (L5) corticothalamic cells. These corticothalamic inputs are formed by the collaterals of the L5b pyramidal tract (PT) cells and arise from all cortical regions studied so far (Bourassa et al., 1995; Harris and Shepherd, 2015; Ojima, 1994). PT cells that send axons to the thalamus also innervate many subcortical sites (Deschenes et al., 1994), thus they represent one of the major pathways via which the cortex can directly impact behavior (Economo et al., 2018; Guillery and Sherman, 2011). In contrast to the L6 corticothalamic axons, the available evidence indicate that the L5 axons do not innervate the TRN (Bourassa et al., 1995; Kakei et al., 2001). Accordingly, the impact of the L5 input on the thalamus is not thought to be sculpted by feedforward inhibition.

In this study using L5-specific transgenic mouse lines we demonstrate that in contrast to other cortical regions L5 PT cells of the frontal cortex innervate the TRN. This suggests a fundamental spatial heterogeneity in corticothalamic communication. Our anatomical and *in vitro* electrophysiology data show qualitative differences between L5-TRN and L6-TRN pathways and our *in vivo* experiments demonstrate that converging L5 activity on TRN neurons is instrumental to determine the correlation between cortical and TRN activity.

## Results

### Selective L5 innervation of the TRN from the frontal cortical areas

To selectively label the axon arbor of the layer 5b (L5) pyramidal cells, floxed AAV-EF1a-DIO-ChR2_EYFP virus was injected to frontal (anterior lateral motor cortex, ALM; primer, M1 and secondary, M2 motor cortices; orbitofrontal cortex, LO, VO; cingulate, Cg and prelimbic cortex, Prl; n=13 injection sites in n=13 mice), and parietal (primer, S1 and secondary S2, somatosensory cortex; n=5 injection sites in n=5 mice) cortical areas of the L5-specific Rbp4-Cre mice (Fig1, A-H). Conditional viral tracing from the parietal regions labelled only passing L5 fibers in the TRN, as described before (Bourassa et al., 1995) (Fig1, A-C). In contrast, frontal cortical areas provided dense L5 collaterals in the anterior part of the TRN studded with boutons (Fig1, D-F). To confirm the presence of the L5-TRN terminals in another mouse line, we mapped the distribution of EYFP-positive terminals in the TRN of the Thy1-ChR2-EYFP mice (Fig1, I), where EYFP is expressed in the L5 cells of the entire neocortex, including regions not targeted by our tracing experiments. Confirming the tracing data, we found labelled Thy1-ChR2-EYFP boutons only in the anterior but not in the posterior TRN sectors. (Fig1, J).

**Figure 1.**
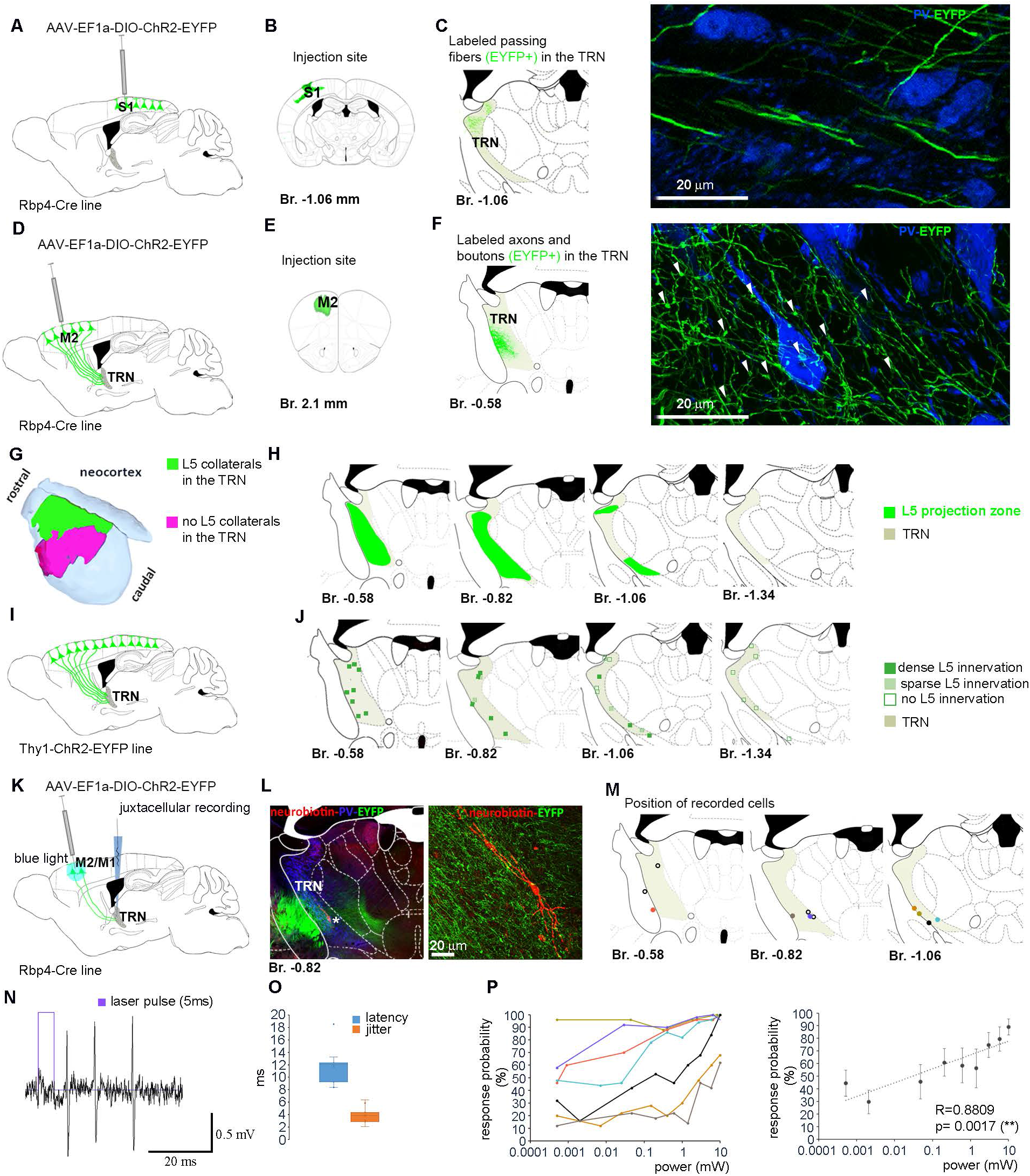
Selective L5 innervation of the TRN from the frontal cortical areas. (A) Experimental design. (B) Schematic figure of the injection site (S1). (C) Schematic figure of the L5 axon distribution in the TRN and high-power confocal image of the L5 axons in the TRN. Note the passing L5 axons without axon terminals. TRN, thalamic reticular nucleus; Br: Bregma. TRN is marked in grey. (D) Experimental design. (E) Schematic figure of the injection site (M2). (F) Schematic figure of the L5 axon distribution in the TRN and high-power confocal image of the L5 axons in the TRN. Arrowheads point to synaptic boutons. Note the dense meshwork of bouton-bearing collaterals. TRN is marked in grey. (G) Cortical regions with (summated areas of n=13 cortical injection sites, green) and without (summated area of n=5 cortical injection sites, magenta) bouton-bearing L5 collaterals. (H) Distribution of L5 collaterals within the TRN. Summated area (green) after n=13 cortical injection sites (see C) at four antero-posterior levels. L5 projection is restricted to the anterior TRN areas. TRN is marked in grey. (I) Experimental design. (J) Distribution of the L5 boutons in the TRN (n=3 mice) in the Thy1-ChR2-EYFP line at four antero-posterior levels. Squares mark the ROI-s of single confocal images depicting dense (dark green), sparse (light green) or no (empty squares) L5 innervation. Note close correspondence with D. (K) Experimental design. (L) Low (left) and high (right) power confocal images of a juxtacellularly recorded and labelled TRN neuron (red) optogenetically activated from the M2 cortex. The neuron is surrounded by a dense meshwork of L5 collaterals (green). Blue in the left image indicates parvalbumin (PV) immunostaining. (M) Position of the cell bodies of the recorded TRN neurons. Colors of the dots match the colors in (P). Cells with a few or only 1 activation intensities (black empty squares) are not involved in (P). (N) Raw data of a single laser stimulus (purple) followed by a TRN response. (O) Box plot for the latency (blue) and jitter (red) of TRN responses at 10 mW laser power value to L5 cortical stimulation (n=11 cells, for each cell average of 50 stimuli). (P) Left panel: Response probabilities of single TRN neurons to L5 stimulation with increasing laser power. (n=7 cells, for each cell 50 stimuli/power value, for the position of the cells see J). Laser power is shown on logarithmic scale. Right panel: Population average for response probabilities at increasing laser power (n=7 cells). Laser power is shown on logarithmic scale. Br: Bregma; TRN: thalamic reticular nucleus

To test whether L5 boutons in the anterior TRN form functional connections we optogenetically activated M1 and M2 L5 cells from the cortical surface (5×10 pulses, 5 ms, 10 mW, at 1 Hz) in anesthetized (ketamin-xylazine) Rbp4-Cre mice injected with AAV-EF1a-DIO-ChR2-EYFP virus. We recorded the evoked responses of TRN cells (n=11 cells in n=6 mice) using the juxtacellular recording and labeling method (Fig1, K-M)(Pinault, 1996). All TRN cells recovered post hoc (Fig1, M, n=11) were located in the anterior TRN. The TRN cells responded with short latency (11.83±0.84 ms, jitter 3.96±0.39 ms) and high fidelity action potentials (response probability: 89.4±6.4 %) (Fig1, N-O). These data confirm functional monosynaptic connection between the cortical L5 and the anterior TRN.

The recruitment of the anterior TRN cells by the L5 pathway was gradual. We tested the response probabilities of both L5 (n=11) and TRN (n=7) cells using different stimulation intensities in the cortex. The number of recruited L5 neurons increased with increasing laser power (FigS1, A-B, Pearson correlation: R=0.9545, ***p=0.0001). While different L5 cells reached threshold at various laser intensities they displayed all-or-none response probability curves (FigS1, A). In contrast, the response probabilities of the TRN cells increased gradually with increasing cortical stimulation power, and the response probability displayed significant correlation with the laser power on a logarithmic scale (Pearson correlation: R=0.8809, **p=0.0017) (Fig1, P). These data together demonstrate the presence of a powerful and selective innervation of the anterior TRN from the L5 of the frontal cortex and imply the convergence of multiple L5 cells on a single TRN cell.

### L5 terminals in the TRN are formed by branching collaterals of corticofugal L5 cells

To test if frontal L5 cells innervating the anterior TRN belong to a specific pyramidal cell population or these cells have other subcortical targets as well, like classical pyramidal tract (PT) neurons, we used the retro-anterograde tracing method (Mátyás et al., 2018). We injected AAV-EF1a-DIO-ChR2-EYFP virus to the upper brainstem of the Rbp4-Cre mice (n=3 mice) (Fig2, A, B) and allowed the mice to survive for 2-3 months. Using this method, the virus first spread retrogradely selectively in the brainstem projecting Rbp4-Cre positive L5 cells, and then its product was transferred to their axonal collaterals in an anterograde manner (Fig2, B). Using this method in all 3 cases, brainstem-projecting L5 cells displayed EYFP-positive axon terminals in the anterior TRN. These terminals were positive for vesicular glutamate transporter type1 (Vglut1) confirming their cortical origin (Fig2, B).

**Fig 2.**
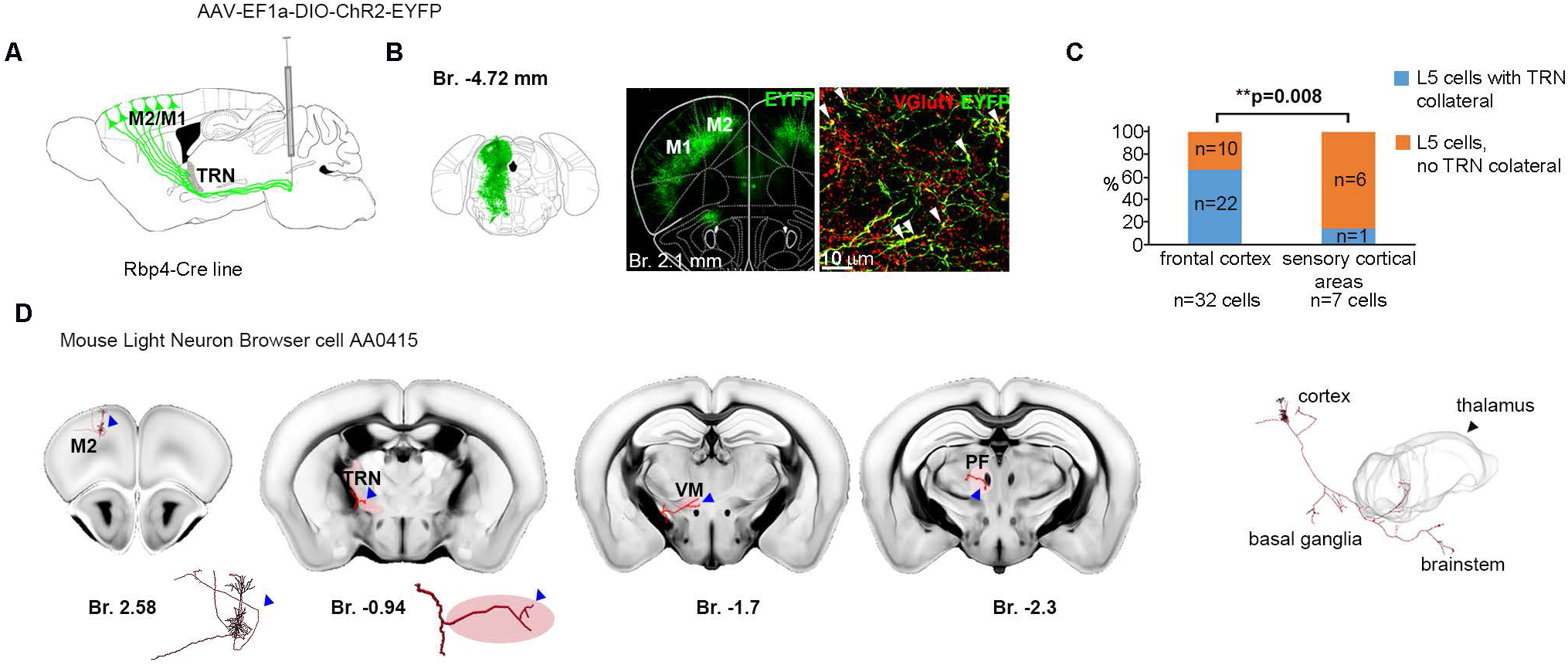
L5 collaterals in the TRN are branching collaterals of the corticofugal L5 cells projecting to the relay nuclei and the brainstem. (A) Experimental design of retro-anterograde tracing. (B) Left panel: injection site in the brainstem. Middle panel: fluorescent image of retrogradely labelled L5 cells in the frontal cortex. Right panel: high magnification confocal image of VGlut1-positive cortical L5 terminals in the anterior TRN. (C) Proportion of corticothalamic L5 cells in the frontal vs. sensory cortices with and without TRN collaterals based on reconstructed L5 cells from the Mouse Light Neuron Browser database. (n=32 cells from the frontal cortex and n=7 cells from the sensory –somatosensory, visual, auditory-cortices). (D) An example L5 cell from the Mouse Light Neuron Browser at four antero-posterior levels. Left panel: position of the soma in the frontal cortex (islet: larger magnification of the cell body). Middle panels: collaterals in the anterior TRN (islet: larger magnification of the L5 collateral) and the VM and Pf thalamic nuclei. Right panel: the complete reconstructed cell with its subcortical targets. Br: Bregma; Pf: parafascicular nucleus; TRN: thalamic reticular nucleus; VM: ventral medial nucleus

To test the presence of L5-TRN collaterals at single cell level, we analyzed the cell reconstruction data of the Mouse Light Neuron Browser database (Chandrashekar, 2017) (39 corticothalamic L5 cells),(TableS1). 22 of the 32 (68.75%) thalamic projecting L5 cells from the frontal cortex emitted collaterals to the anterior TRN (Fig2, C-D), demonstrating high frequency of L5-TRN collaterals in this population. In case of L5 cells from the sensory cortices, only 1 out of 7 (14.29%) had TRN collateral (Chi-square test: **p=0.008) (Fig2, C). All the TRN projecting cells sent axons to the basal ganglia and the brainstem as well (Fig2, D). These two datasets together confirm that TRN projecting L5 cells belong to the classical brainstem projecting PT cells.

### Parallel, but morphologically distinct L5- vs. L6-TRN pathways

Frontal cortical areas are known to target the TRN via L6 pyramidal cells (Jones, 1975; Lozsadi, 1994). Thus, we examined whether L6 and L5 inputs from the same cortical area form parallel or divergent pathways in the TRN and whether they are morphologically AND/OR functionally different. To simultaneously label both pathways, we injected a mixture of CreON and CreOFF viruses to the M2 of the L6-specific Ntsr1-Cre mouse (n=3 mice) (Fig3, A). At the injection site the AAV-EF1a-DIO-ChR2-mCherry virus (CreON, red) was expressed selectively in the L6 corticothalamic cells in a Cre-dependent manner whereas the AAV-DFO-ChR2-eYFP virus (CreOFF, green) was expressed exclusively in the cortical cells that did not contain the Cre-recombinase (including the L5 corticothalamic cells). Since thalamus receives cortical inputs only from L6 and L5 corticothalamic cells (Rouiller and Welker, 2000), we could reliably label the two populations in parallel within the same animal using this method (Fig3, B). In all 3 animals, L6 and L5 projection zones displayed strong overlap in the anterior TRN (Fig3, C) implying that L6 and L5 inputs from the same cortical source converge in the same TRN zone.

**Figure 3.**
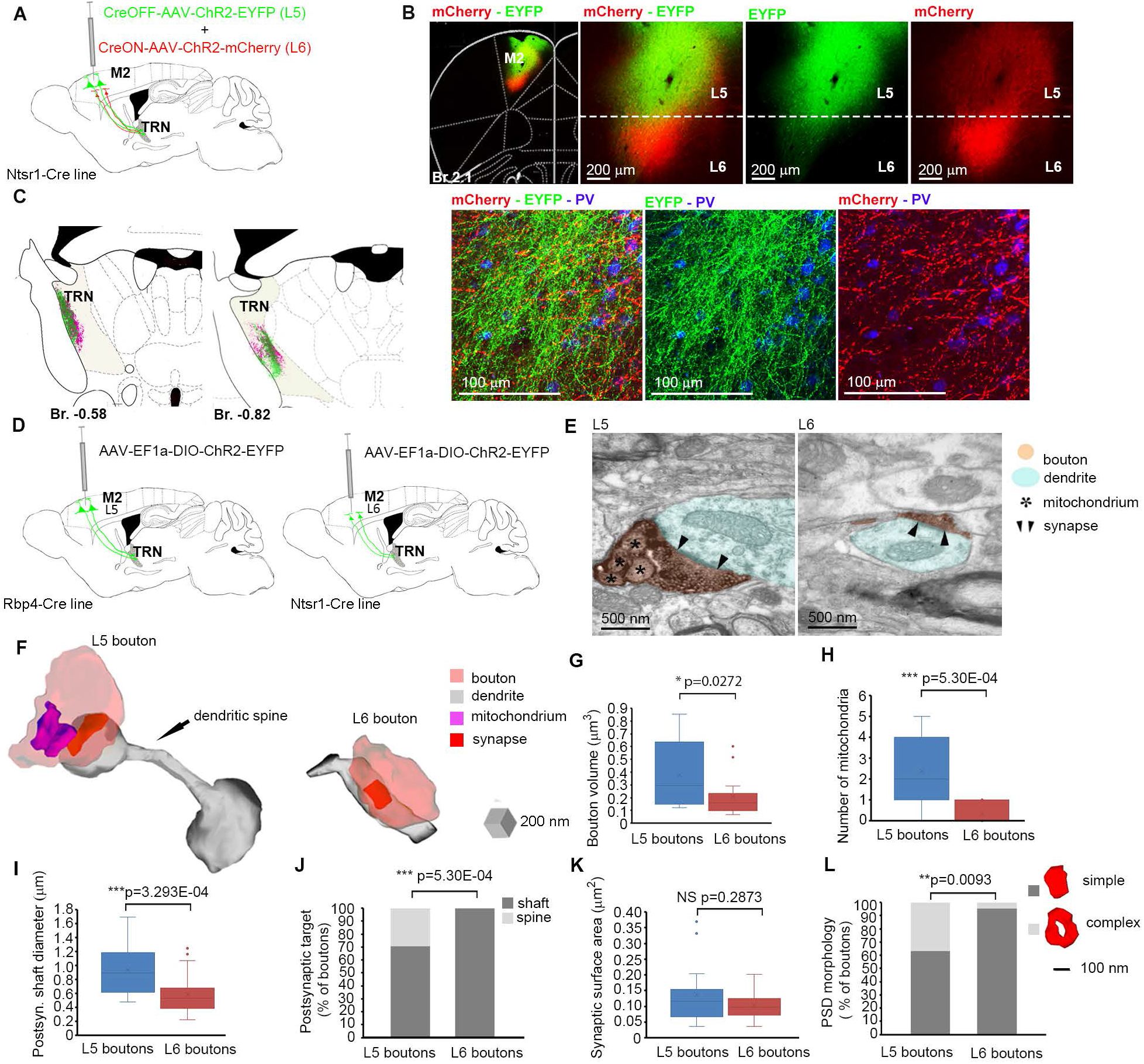
Parallel, but distinct L5- vs. L6-TRN pathways. (A) Experimental design to simultaneously label L5 and L6 projections in the L6 specific Ntsr1-Cre line. (B) Low power (left) and high power (middle left) fluorescent images of the injection site (mCherry, red, Ntsr1-positive cells; EYFP, green, Ntsr1-negative cells) in the M2 cortex. Middle right and right panels, separate green and red channels depicting non-overlapping expression of the two markers. (C) Schematic drawing depicting overlapping L6 (red) and L5 (green) projection zones in the TRN at two antero-posterior levels (left two panels). Right three panels, high magnification confocal images of the L6 and L5 projection zones in the TRN (PV+, blue). (D) Experimental design. (E) High power electron micrographs showing examples for a L5 (left panel) and a L6 (right panel) terminal (black precipitate, orange shading) in the TRN establishing synapses on a middle and small calibre dendrite (blue shading), respectively. Edges of postsynaptic densities are marked by black arrowheads; mitochondria are marked by black asterisks. (F) Examples for 3D-EM reconstructed L5 (left) and L6 (right) boutons and their postsynaptic targets in the TRN. The L5 bouton targets a dendritic spine marked by black arrow, the L6 bouton synapses on a thin dendritic shaft. Orange, bouton; grey, postsynaptic target; red, synaptic surface; purple, mitochondrium. (G) Boxplots for the volumes of L5 (blue, n=14) and L6 (red, n=13) boutons. (H) Boxplots for number of mitochondria in the L5 (blue, n=15) and L6 (red, n=15) boutons. (I) Boxplots for the diameter of postsynaptic dendritic shafts for L5 (blue, n=12) vs. L6 terminals (red, n=40) in the TRN. (J) Postsynaptic targets of L5 (n=17) vs. L6 (n=40) terminals in the TRN. (K) Boxplots for the synaptic surface area of the L5 (blue, n=19) and L6 (red, n=24) synapses in the TRN. (L) Morphological types of the postsynaptic density (PSD) in the L5 (n=19) and L6 (n=22) synapse populations in the TRN. Br: Bregma; TRN: thalamic reticular nucleus

The functional properties of a synapse are closely related to the ultrastructure of its pre- and postsynaptic elements. Thus, we compared L6 and L5 synapses at electron microscopic (EM) level. L6 and L5 cells were virally labeled in the frontal cortex of the Ntsr1-Cre and Rbp4-Cre mouse line respectively (Fig3, D). Both L5 and L6 terminals established classical asymmetrical synapses in the TRN (Fig3, E). Serial EM sections of the labelled L6 and L5 synapses were analyzed, and the boutons and their postsynaptic partners were reconstructed in 3D (Fig3, F). L5 boutons had significantly larger volume (L5: n=14, 0.38±0.07 µm^3^; L6: n=13 0.2±0.05 µm^3^; Mann-Whitney U Test: *p=0.0272), (Fig3, G), and contained, in most of the cases, multiple mitochondria. L6 boutons, on the other hand, contained no or maximum 1 mitochondrium (L5: n=15, 2±0.46; L6: n=15, 0.33±0.13, Mann-Whitney U Test: ***p=5.3E-04), (Fig3, H). L5 boutons targeted significantly thicker dendrites compared to the L6 boutons, suggesting that they may prefer different dendritic compartments (L5: n=12, 0.94±0.1 µm5; L6: n=40; 0.59±0.04 µm, Mann-Whitney U Test: **p=0.0016) (Fig3, I). In line with this 29.41% of the L5 boutons (n=17) targeted dendritic spines, whereas L6 (n=40) boutons targeted exclusively dendritic shafts (Chi-square test: ***p=3.293E-04) (Fig3, J). Although there was no significant difference between the synaptic surface area of the L5 and L6 synapses (L5: n=19, n=24 0.14±0.02 µm^2^, L6:0.1±0.01 µm^2^, Mann-Whitney U Test: NS p=0.2873) (Fig3, K), 36.84% of the L5 synapses (n=19) had complex (perforated or branching) morphology, whereas L6 synapses (n=22) showed in all but one case (4.55%) simple, discoid morphology (Chi-square test: **p=0.0093) (Fig3, L). These data shows that the L6-TRN and L5-TRN pathways have different ultrastructural characteristics both regarding the pre- and the postsynaptic elements suggesting distinct functional properties.

### Distinct physiological properties of the L5-TRN and the L6-TRN pathways

To compare the synaptic properties of L5-TRN and L6-TRN pathways we used *in vitro* electrophysiology experiments. AAV-EF1a-DIO-ChR2-EYFP virus was injected 2-4 weeks before the experiments in the frontal cortex of the Ntsr1-Cre or Rbp4-Cre mouse to label L6 and L5 pathways respectively. L5 and L6 fibers were optogenetically activated while TRN cells were recorded in whole-cell patch clamp configuration (Fig4, A-B). TRN neurons in Rbp4-Cre and Ntsr1-Cre mice showed similar passive cellular properties (Rbp4-Cre: n = 18 neurons, Ntsr1-Cre: n = 10 neurons; membrane resistance (Rm): 287±30 MΩ vs. 264±25MΩ, membrane capacitance (Cm): 81±8 pF vs. 58±8 pF, resting membrane potential (RMP): 59±2 mV vs. −64±4 mV; Student’s t tests, p = 0.56 for Rm, *p= 0.042 for Cm and p = 0.29 for RMP) (Fig4, C). Evoked synaptic currents of both pathways were of overall small mean amplitude even for maximal levels (3.5 mW, 1 ms) of light stimulation (L5-TRN: n=15 neurons, −15 to −242 pA, mean 74 ± 13 pA; L6-TRN: n=10 neurons −4 to −77 pA, mean −25 ± 8 pA). The rise and decay times of the L5-TRN EPSCs were significantly longer compared to the L6-TRN EPSCs (L5-TRN: n=15 neurons, L6-TRN: n=9 neurons; rise time: 0.8±0.09 ms vs. 0.43±0.04 ms; decay time: 4.67±0.76 vs. 2.87±0.30 ms; Student’s t test, ***p = 9E-04 for rise time, *p = 0.042 for decay time). The response latency was shorter in case of the L5-TRN pathway (2.62±0.12 vs. 3.86±0.45 ms, *p = 0.026) (Fig4, D). The L5-TRN EPSCs had significantly higher NMDA-receptor-mediated component (n = 4 for L5-TRN and n = 4 for L6-TRN synapses; 20±3% vs. 8±1%; Student’s t test, *p = 0.02) (Fig4, E).

**Figure 4.**
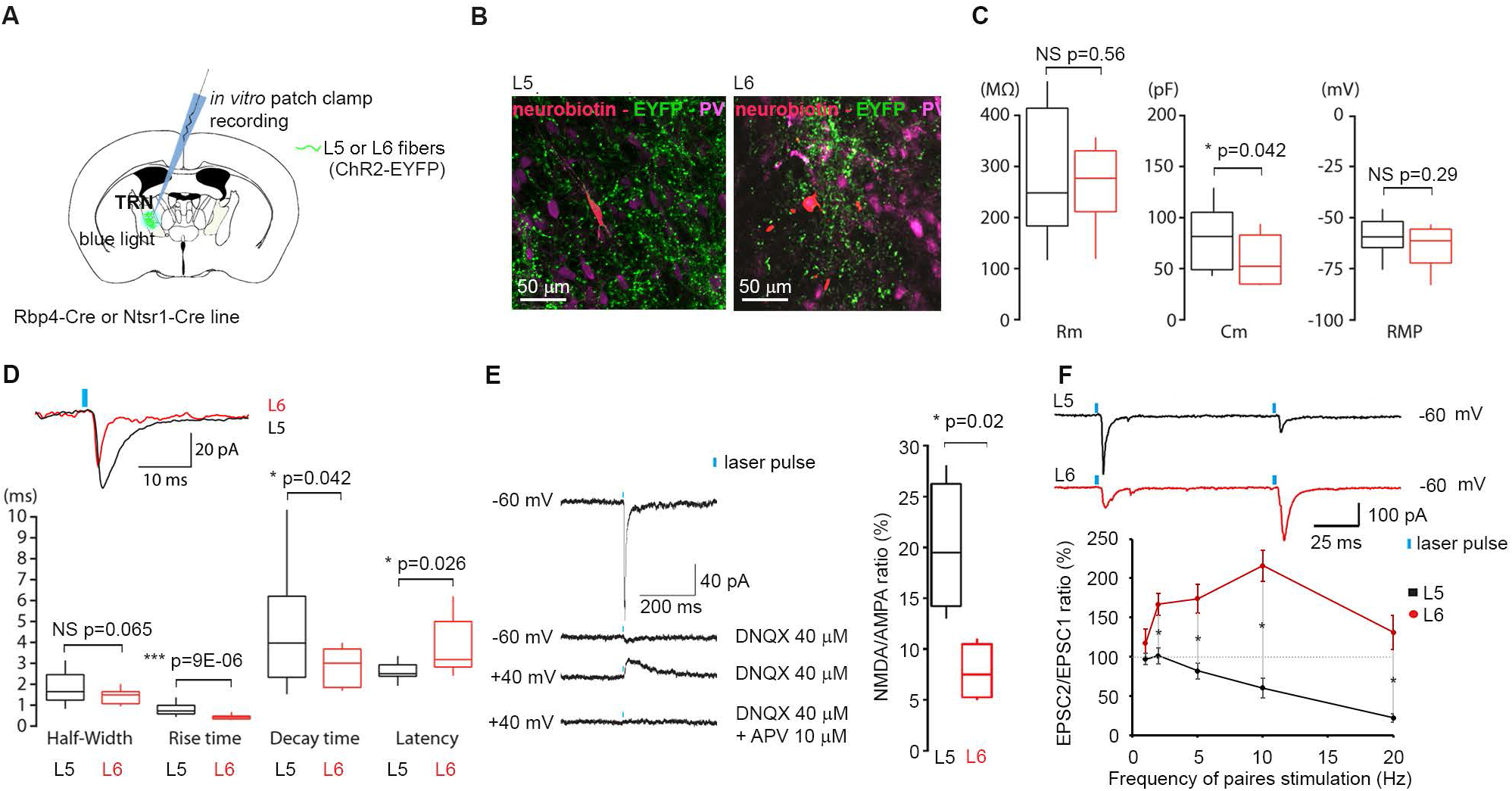
Distinct physiological properties of the L5-TRN and the L6-TRN pathways. (A) Experimental design. (B) Neurobiotin labelled TRN cells (red) in the PV-positive TRN (magenta) among L5 (left panel) or L6 (right panel) fibres (ChR2-EYFP+, green). (C) Box-and-whisker plots showing the membrane resistance (Rm), the membrane capacitance (Cm) and the resting membrane potential (RMP) of the recorded TRN neurons. TRN neurons in Rbp4-Cre (L5, black) and Ntsr1-Cre (L6, red) mice showed similar passive cellular properties (Student’s t tests, n = 18 for Rbp4 and n = 10 for Ntsr1). (D) Box-and-whisker plot showing the evoked postsynaptic currents half-width, rise time, decay time and latency from LED onset for L5-TRN (black) and L6-TRN synapses (red). (Student’s t test, n = 15 for L5-TRN and n = 9 for L6-TRN) Islet: Examples for L5 (black) and L6 (red) EPSCs. (E) left panel: Typical traces of NMDA/AMPA receptor-mediated synaptic responses evoked by optogenetic activation of L5 axons in TRN. Right panel: Boxplots of the NMDA/AMPA ratio in case of L5-TRN (black) and L6-TRN (red) synapses (Student’s t test, n = 4 for L5-TRN and n = 4 for L6-TRN). (F) Top: typical traces from voltage-clamp recording of TRN neurons upon paired light activation of L5 (black) and L6 (red) afferents from the motor cortex. Bottom: graph of the paired-pulse ratio of evoked postsynaptic currents (EPSCs) in TRN neurons at 1, 2, 5, 10 and 20 Hz (n = 7 for L5 -TRN and n = 10 for L6-TRN; Wilcoxon rank sum test with Bonferroni correction for multiple comparison).

The short-term plasticity of the L5-TRN and L6-TRN pathways displayed opposite features. While paired-pulse activation of L5 afferents showed clear short-term depression, L6 activation showed typical short-term facilitation described previously for L6 projections in somatosensory cortex (Astori and Lüthi, 2013; Fernandez et al., 2018), (n = 18 cells for L5-TRN and n = 10 cells for L6-TRN synapses; Wilcoxon rank sum test with Bonferroni correction for multiple comparison (significance at 0.01)) (Fig4, F).

In summary the L5-TRN synapses showed higher NMDAR content than L6-TRN synapses, which might explain differences in the EPSC time course, notably its decay time. Moreover, the two pathways showed different forms of short-term plasticity. Strong short-term depression of the L5-TRN pathway implies that it may be better tuned for the integration of instantaneous and synchronous L5 activity than to faithfully follow long spike trains.

### Segregation and integration of different L5 inputs in the TRN

The previous *in vivo* electrophysiology experiments (Fig1, K-P, FigS1, A-B) implied the convergence of multiple L5 cells on a single TRN cell. Therefore, we asked to what extent the frontal cortical L5 activity could be integrated at the level of single TRN cells. To address this question, double viral injections (AAV-EF1a-DIO-ChR2-EYFP and AAV-EF1a-DIO-ChR2-mCherry) with non-overlapping injection sites were made to various combinations of frontal cortical regions in the Rbp4-Cre mice (Fig5, A-B). In all cases, we saw clear segregation of L5 collaterals arising from neighboring cortical regions in the TRN (Fig5, C-D). Using the results of multiple double and single viral labelling experiments (M2 rostral + M2 caudal (n=1); M2 rostral + M1 (n=1); M2 rostral + mPFC rostal (n=1); mPFC rostral (n=1); mPFC caudal (n=2), orbitofrontal cortex (n=2)), we created the map of the L5-TRN pathway (Fig5, E). We found segregated, patchy organization of L5-TRN termination zones. As an indication for a topographical organization, we found that within one cortical region, axons from more caudal cortical areas targeted preferentially more dorsal parts of the TRN compared to the axons from the more rostral areas. We compared these tracing data with single cell results of the Mouse Light Neuron Browser database. Areal localization of the corticothalamic (n=22) cells and the position of their TRN collaterals (n=26) were consistent with our viral tracing data (Fig5, F-G).

**Figure 5.**
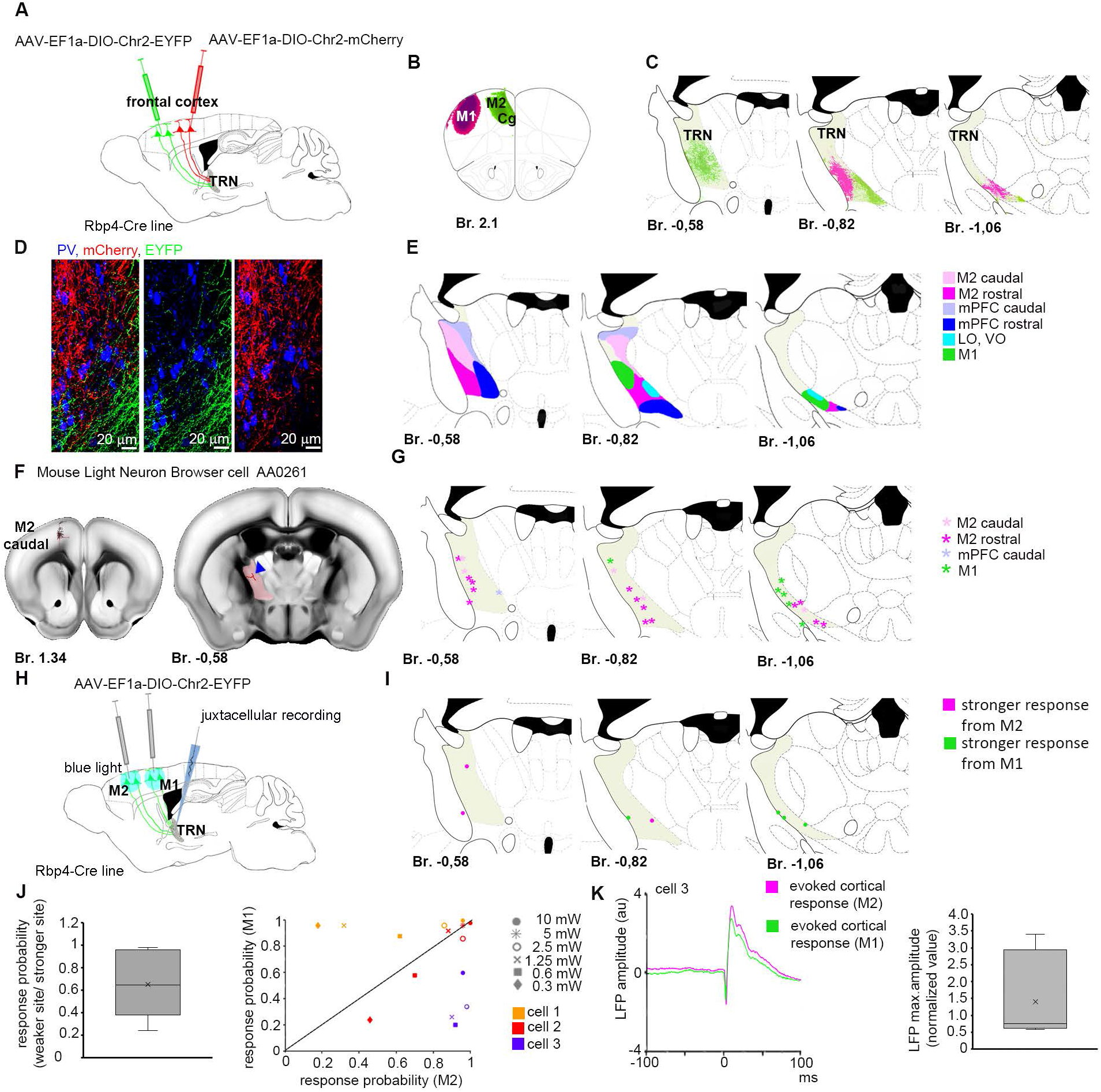
Topography of the L5-TRN pathway. (A) Experimental design. (B) Schematic drawing of non-overlapping injection sites in the M1 (magenta) and M2/Cg (green) cortices. (C) Schematic drawing of the non-overlapping L5 projection zones in the TRN following injections shown in B at three antero-posterior levels. (D) High power confocal images of the non-overlapping M1 (red) and M2/Cg L5 (green) collaterals in the TRN (PV+, blue). Same mice as (B) and (C). (E) Topographical map of the frontal L5 to anterior TRN projection (n=11 injections of which 6 dual and 5 single injections). TRN is labelled in grey. (F) A reconstructed L5 corticothalamic cell from the Mouse Light Neuron Browser. Left panel: Position of the cell body in the M2. Right panel: position of axon collateral (red) in the TRN (labelled with pale red). (G) Position of the axon collaterals in the TRN for n=22 reconstructed L5 corticothalamic cells from the Mouse Light Neuron Browser. Some neurons had collaterals at more than one TRN levels. Colors indicate the position of the L5 soma. Rostral and caudal levels are defined at the Br. 1.8 level to match the nomenclature of our double injection experiments in (E). TRN is labelled in grey. Note close correspondence between the track tracing (E) and single cell labelling data. (H) Experimental design for dual optogenetic activation. (I) Position of the cell bodies of the recorded TRN cells. Colors indicate the site of cortical activation with stronger TRN responses. (J) Left panel: Box plot depicting the ratio of TRN response probabilities from cortical stimulation sites with the lower vs higher TRN responses normalized to the response probabilities from the stronger site. (n=8 cells in n=5 mice). Right panel: Comparison of the response probabilities after M1 vs. M2 stimulation in three TRN cells at different laser intensities. Symbols indicate laser powers, colors label different cells. Note different preference for stimulation sites. (K) Left panel: An example for evoked cortical response averages at the sites with the weaker and with the stronger TRN responses. (L) Right panel: Box plot for the peak amplitudes of the evoked cortical response averages at the sites with the weaker TRN response probabilities normalized to the response peak amplitudes at the stronger cortical sites. (n=8 cells in n=5 mice). au: arbitrary unit Br: Bregma; Cg: cingulate cortex; LO: lateral orbital cortex; mPFC: medial prefrontal cortex; TRN: thalamic reticular nucleus; VO: ventral orbital cortex

Next, we tested whether single TRN neurons are able to integrate inputs from two different cortical regions by stimulating L5 cells in the M1 and M2 areas (same rostrocaudal, but different mediolateral level) in the Rbp4-Cre mouse (n=7 cells), (Fig5, H-I). All TRN neurons had evoked responses from both cortical regions. The preferred stimulation site for each cell was determined as the site with higher response probability (P) at 10 mW stimulus power (n=7 cells in n=5 mice; P weaker/stronger site = 0.63±0.09) (Fig5, J). We found cells both with strong M1 or M2 preference or with comparable response probabilities for the two stimulation sites (Fig5, J). Matching our viral tracing data (Fig5, E), somata of TRN cells with stronger response to M1 stimulation were positioned in more caudal TRN regions, compared to the TRN cells that preferred M2 stimulation (Fig5, I). Evoked cortical LFP responses were of comparable magnitude at the sites evoking strong or weak responses in TRN (evoked response peak weaker/stronger site = 1.41±0.35), showing that the preference of the TRN cells was not due to uneven activation of cortical sites (Fig5, K).

These data together demonstrate that that L5 inputs from different cortical regions segregate at the level of the TRN. However, individual TRN cells are able to integrate inputs from different cortical regions probably via their extensive dendritic arbors spanning multiple cortical termination zones.

### Gradual recruitment of L5 pyramidal cell is reflected in TRN spike output

Relatively weak L5-mediated synaptic responses *in vitro* (Fig4, D, F) could elicit reliable responses in the TRN *in vivo* (Fig1, K-P) often from broad cortical origins (Fig 5, H-K). This suggests a significant amount of convergence in the L5-TRN pathway. This was further supported by the gradual recruitment of TRN cells with increasing laser intensities (Fig1, P) in contrast to the all-or-none responses of L5 cells (FigS1, A-B). This implies that convergent, individually weak L5 inputs might allow TRN cells to faithfully read out gradual changes in cortical population activity.

To test this idea, we optogenetically increased cortical L5 population activity step by step and examined the properties of the evoked TRN responses (n=20 cells in n=9 mice) in the Thy1-ChR2-EYFP mice (Fig6, A-E). Evoked responses ranged from single spikes, to bursts with up to 564 Hz average intraburst frequencies (aIBF) (Fig6, D; FigS1 C-D). Individual bursts contained 2-16 spikes (FigS1, E-F). Single spike events and bursts with broad range of aIBFs (100-369 Hz), (Fig6, D; FigS1, G-H) could also be observed during the spontaneous activity of the same cells. aIBFs of 77.29 % of the evoked bursts were in the range of the values observed during baseline activity (FigS1, H). This indicates that the majority of the optogenetically evoked activity was within the physiological range.

**Figure 6.**
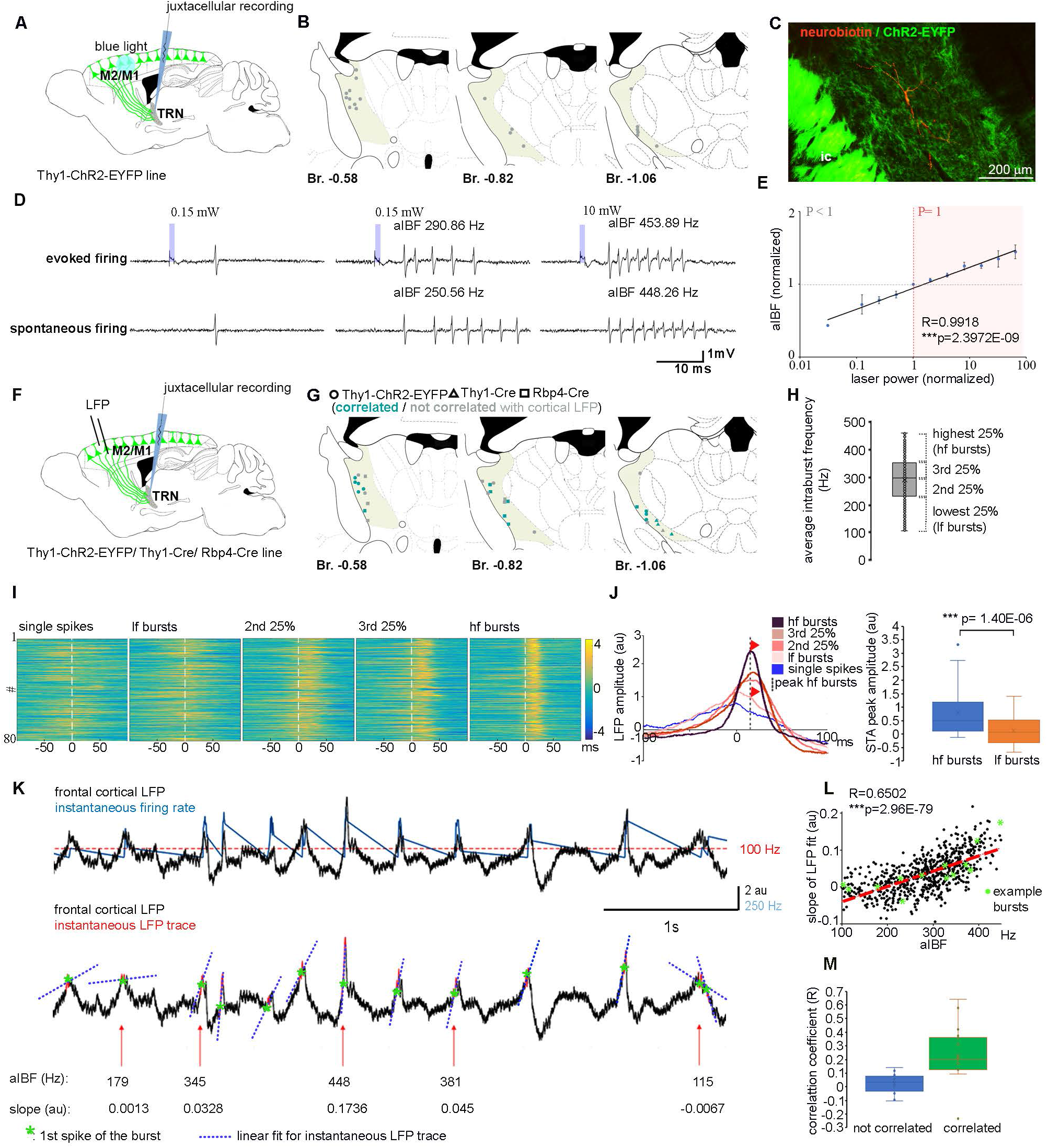
Instantaneous correlation between TRN firing and cortical LFP activity. (A) Experimental design. (B) Position of juxtacellularly recorded and filled TRN cells. (C) Confocal image of a recorded and neurobiotin filled TRN cell (red) surrounded by ChR2-EYFP positive L5 fibers (green). (D) Examples of evoked and spontaneous TRN single spike events and bursts. blue: optogenetic activation. Power of the optogenetic stimulus is indicated above. aIBF: average intraburst frequency. (E) Normalized aIBF averages for n=20 cells in n=9 mice shown in a log-linear scale. Both the aIBFs and the laser power values are normalized to the value observed at the threshold power. Threshold power was defined as the minimal laser power value at which P=1 response probability was observed. aIBFs showed significant log-linear correlation with the laser power. (F) Experimental design. (G) Position of juxtacellularly recorded and filled TRN cells. Circles, triangles and squares label cells from the Thy1-ChR2-EYFP Thy1-Cre and Rbp4-Cre lines respectively. (H) aIBFs of spontaneous bursts for a TRN cell. Same as (F). aIBFs were grouped into four quartiles with increasing values (lf. 2^nd^ 25%, 3^rd^ 25%, hf). (I) Color plots of instantaneous LFP traces for single TRN spikes and bursts grouped into four quartiles as shown in (H) for an example cell. Colors indicate the amplitude values of the standardized LFP traces. Rows represent single TRN spike or burst events. 0 marks the peak of the single spike or of the first spike in a burst. For the better comparison, equal number of events for all firing categories were selected randomly. (J) Left panel: Average LFP traces (spike triggered averages, STAs) for each spiking category for the same cell shown in (L). Dashed line, STA peak for high frequency (hf) bursts. Because STA peaks for the low frequency (lf) bursts were often less well defined, we calculated the STA amplitude for lf bursts at the position of the hf STA peak (read arrowhead). Note progressively higher amplitude and narrower STA for higher frequency bursts. Right panel: Box plots of the population averages (n=31 cells in n=13 mice) for the peak STA values of lf and hf TRN bursts (Student’s paired sample t-test). (K) Upper panel: A raw frontal cortical LFP trace (black) and the concurrent instantaneous firing rate (blue) of a TRN cell. Red dashed line indicates burst threshold (100 Hz). Lower panel: Calculation of the linear fit for the instantaneous LFP trace on the same LFP trace (see Methods). Green asterisk, first spikes of the TRN bursts. Red, instantaneous LFP trace (30ms long LFP sections, timed to the first spike of the TRN burst). Blue dashed line, linear fits for these 30ms sections. aIBFs of the TRN cell and the slope values for the linear LFP fits are indicated for each burst. (L) Correlation between the aIBFs of the bursts and the slope of the instantaneous LFP traces for n=649 spontaneous burst of the example TRN neuron shown in N (Pearson correlation). Green asterisks: example bursts on (N). (M) Right panel: Box plots for the Pearson correlation coefficients for aIBF - LFP slope correlations for n=31 cells. Green, cells with significant aIBF – LFP slope correlation (n=19 cells in n=11 mice); blue: cells with no significant correlation (n=12 cells in n=7 mice). au: arbitrary unit; ic: internal capsule

Both the aIBFs and the number of spikes per evoked bursts showed significant log-linear correlation with the cortical L5 stimulus power (Pearson correlation: R=0.9918, ***p=2.3972E-09 and R=0.9702, ***p=7.6329E-07 respectively) (Fig6, E; FigS1, C-F). Since the fraction of recruited L5 cells showed similar, log-linear correlation with the laser power (FigS1 A-B), these data demonstrate that the exact spike output of the TRN cells reflects the number of simultaneously recruited L5 cells.

### Instantaneous correlation between spontaneous cortical and TRN activity

To investigate whether TRN spiking was also modulated by spontaneous changes in the cortical synchrony we recorded the spontaneous (baseline) activity of n=44 TRN cells in parallel with the frontal cortical LFP (Fig6, F). Synchronous cortical population activity similar to the optogenetically evoked cortical responses could be detected in the LFP recordings as transient, fast, high amplitude events. Under our conditions, fast LFP transients were present only in light but not in deep anesthesia (FigS2, A-D). In deep anesthesia, cortical LFP activity was dominated by a regular, slow (1-4 Hz) component (2.66±.0.46 dB), (n=13 cells in n=8 mice, from which Thy1-ChR2-EYFP (n=5), Rbp4-Cre (n=3)), (FigS2, B-C). Thus, we used the recordings under light anesthesia for further analysis (see Materials and Methods) (n=31 cells in n= 13 mice, from which Thy1-ChR2-EYFP (n=5), Rbp4-Cre (n=6) and Thy1-Cre (n=2)) (Fig6, G).

Examination of individual TRN spike triggered LFPs, and spike-triggered averages (STA) of LFPs, showed that spontaneous single spikes fired by TRN cells were mostly associated with irregular cortical activity. In contrast, TRN bursts were associated with fast LFP transients (Fig6, H-J; FigS2, D). Bursts with higher aIBFs had a population STA with progressively higher peak amplitude indicating that faster bursts are better synchronized with higher amplitude cortical events compared to slower bursts (Fig6 I-J) (peak STA amplitudes;0.14±0.11 arbitrary unit (au) for the burst with the lowest 25% of aIBFs vs. 0.81±0.14 au for the burst with the highest 25% of aIBFs; Student’s paired sample t-test: - ***p= 1.4 E-06).

To quantify the gradual, instantaneous relationship between cortical population activity and TRN firing we correlated the aIBFs of the individual TRN bursts and the slope of the corresponding cortical LFP transients (see Materials and Methods), (n=31 cells and n= 12531 bursts in n= 13 mice, from which Thy1-ChR2-EYFP (n=5), Rbp4-Cre (n=6) and Thy1-Cre (n=2)). In 19 out of 31 TRN cells (61.3%) there was a significant correlation between the aIBF of the bursts and the magnitude of the instantaneous LFP slope (n=19 cells in n=11 mice, from which Thy1-ChR2-EYFP (n=5), Rbp4-Cre (n=4) and Thy1-Cre (n=2); Pearson correlation: R=0.230±0.051 *p<0.05 n=4 cells; **p<0.01 in n=1 cells; ***p<0.001 in n=14 cells) (Fig6 K-M). TRN bursts with higher aIBFs were correlated with faster LFP events. These data clearly demonstrate a tight link between cortical population activity and the exact spiking output of the majority of TRN cells.

Synchrony of TRN and cortical activity can arise not only from L5 neurons as proposed here but also from the relay cells, which innervate both the TRN and the cortex. To address this question, we recorded the activity of TRN cells (13 TRN cells in n=5 mice), thalamocortical cells in ventromedial relay nucleus, which has frontal cortical connections (VM, n=5 cells in n=5 mice) and L5 cortical neurons (n=6 cells in n=2 mice) in the frontal cortex in the Thy1-ChR2-EYFP line. We compared the activity of these cells during the fast cortical LFP transients (see Materials and Methods). In our sample, 6 out of the 13 TRN cells elevated their firing around the peaks (± 50ms) of the transients (FigS3, A1-D1). Three out of 6 L5 cells also strongly modulated their activity during the fast transients (FigS3, A2-D2). However, we could not find modulated cells in the VM population (FigS3, A3-D3). These data suggest that the elevated cortical, not thalamic, firing underlies the recruitment of TRN bursting at fast cortical transients.

Taken together we found that both in case of the evoked and spontaneous cortical population events the exact spike output of the anterior TRN cells provides a gradual read-out of the magnitude of synchronous cortical activity.

### Heterogeneity of anterior TRN cells

In 12 out of the 31 TRN neurons (38.7%) bursts were either not associated with cortical LFP transients (FigS2, B) or the burst properties were not correlated with the magnitude of the slope of the cortical events (n=12 cells in n=7 mice, from which Thy1-ChR2-EYFP (n=2), Rbp4-Cre (n=4) and Thy1-Cre (n=1); Pearson correlation: R=0.027±0.022, p>0.05 in n=12 cells) (Fig6, M). TRN cells with or without significant aIBF-LFP slope correlation were mixed spatially within the anterior TRN (Fig6, G) and did not differ in aIBFs (233.72±8.9 vs. 170.71±12.44 Hz Mann-Whitney U Test; p= 0.09) (FigS2, H) or spikes per bursts values (5.36±0.34 vs. 3.99±0.50 Hz Mann-Whitney U Test; p= 0.1964) (FigS2, G). Correlated anterior TRN cells, however had significantly higher firing rates (9.17±1.17 vs. 6.06±1.32 Hz Mann-Whitney U Test; p= 1.54E-04) (FigS2, E) and burst rates (0.5±0.03 vs. 0.34±0.06 Hz Mann-Whitney U Test; p= 5.20E-03) (FigS2, F). These data indicate that TRN populations are heterogeneous in the anterior TRN concerning their intrinsic and network properties.

In deep anesthesia we did not find significant aIBF-LFP slope correlation (n=13 cells in n= 11 mice, from which Thy1-ChR2-EYFP (n=5), Rbp4-Cre (n=4) and Thy1-Cre (n=2)), (Pearson correlation: R=0.13±0.06, p>0.05 in n=13 cells) (FigS2, A-D). In these cases both the firing rate (6.29±1.53 Hz vs. 9.17±1.17 Mann-Whitney U Test; p= 3.63E-07) (FigS2, E), the burst rate (0.32±0.07 Hz vs. 0.5±0.03 Mann-Whitney U Test; p= 3.17E-05) (FigS2, F., the number of spikes per burst (3.99±0.50 Hz vs. 5.36±0.34 Mann-Whitney U Test; p= 0.0262) (FigS2, G) and the aIBF of the bursts (170±12.44 Hz vs. 223.72±8.90 Mann-Whitney U Test; p=0.0017) (FigS2, H) were significantly lower compared to the cells with significant aIBF-LFP slope correlation in light anesthesia.

### Perturbation of the L5 signaling in the anterior TRN diminishes the instantaneous correlation between TRN firing and cortical LFP activity

To test if L5 input is necessary for recruiting TRN neurons during cortical fast LFP transients we selectively perturbed the activity of L5 terminals in the anterior TRN by optogenetically activating the ArchT inhibitory opsin in the Rbp4-Cre and Thy1-Cre transgenic line (Fig7, A). We selected TRN cells (Fig7, B) for further recording by applying 2-3 test pulses (5s each) of yellow laser, which caused a transient, slight decrease in the firing rate (see Materials and Methods). For data analysis only cells with significant positive baseline aIBF-LFP slope correlation were used (n=7 cells in n=4 mice from which n=2 Rbp4-Cre and n=2 Thy1-Cre mice). Position of the recorded cells, and the optic fiber tip and the topography of labelled L5 fibers were verified post hoc (Fig7, C). After recording the baseline activity of the cells, we applied yellow light (5s ON 10s OFF, 60 cycles) to perturb the L5 terminals. Upon optogenetic activation of ArchT, we did not observe a persistent alteration in the firing rate (6.89±1.76 Hz vs. 6.65±1.91 Hz Student’s paired sample t-test: p= 0.8208) or burst rate of TRN cells (0.59±0.04 Hz vs. 0.58±0.06 Hz Student’s paired sample t-test: p= 0.8279) (Fig7, D; FigS4, A and B). Similarly, the aIBFs (227.25±16.63 Hz vs. 225.49±19.59 Hz; Student’s paired sample t-test: p= 0.7962) and number of spikes per bursts (5.71±0.39 vs. 5.27±0.44; Student’s paired sample t-test: p= 0.1389) were unaffected by the manipulation (Fig7, D; FigS4, C-D).

**Figure 7.**
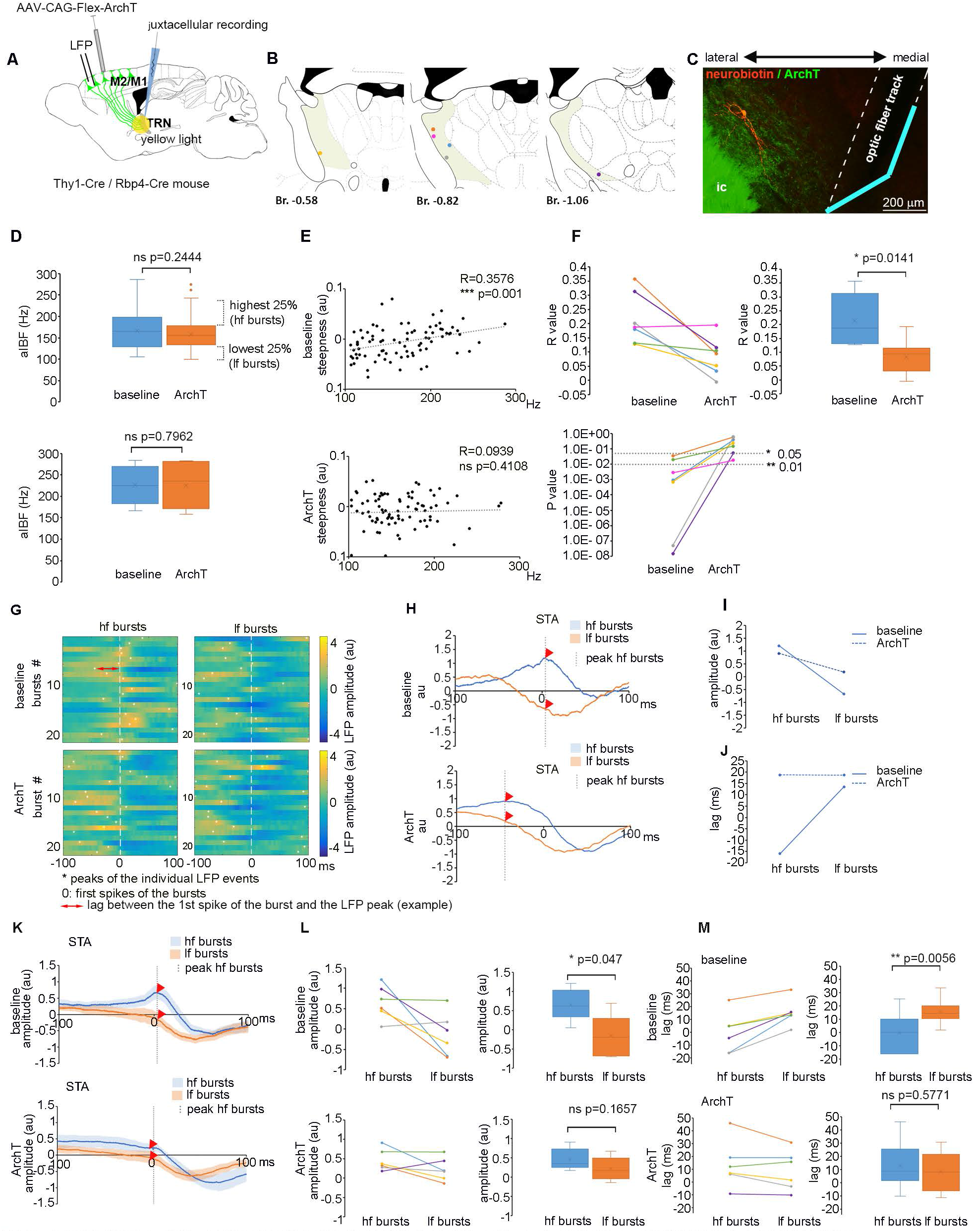
Perturbation of the L5 input in the anterior TRN diminishes the instantaneous correlation between TRN firing and cortical LFP activity. (A) Schematics of experimental design to selectively perturb L5 inputs in the anterior TRN, and simultaneously record TRN activity and frontal cortical LFP. (B) Position of juxtacellularly recorded and filled TRN cells. Colors of the dots match colors in (F), (L) and (M). (C) Confocal image of a neurobiotin filled TRN cell (red) surrounded by ArchT-EYFP positive L5 fibers (green). The position of the optic fiber track is indicated by white dashed line. Blue line labels the orientation of the mirror in the optic fiber tip. (D) upper panel: Box plots of the aIBF of TRN bursts during baseline activity (blue) and during optogenetic perturbation (ArchT, orange) of L5 inputs for a TRN cell (baseline: n=83 bursts; ArchT: n=313 bursts) (Mann-Whitney U Test) lower panel: Same as upper panel for a population of n=7 TRN cells (Student’s paired sample t-test). (E) Correlation of the aIBF of the bursts and the slope of the instantaneous cortical LFP trace for an example TRN cell. upper panel: baseline activity (Pearson correlation); lower panel: optogenetical perturbation of L5 inputs (Pearson correlation); To calculate R and P values, equal number of bursts were randomly selected via multiple resampling for each condition (see Materials and Methods). (F) upper left panel: Pearson correlation coefficients (R values) for the aIBF - LFP slope correlation for (n=7 cells, shown with different colors) at baseline conditions and during optogenetic perturbation of L5 inputs. upper right panel: Box plots for the values shown in the upper left panel. (Student’s paired sample t-test). lower panel: P values of Pearson correlation for n=7 TRN cells at baseline vs. during optogenetical perturbation of L5 inputs (ArchT). 1% and 5% significance levels are indicated with dashed lines. (G) Color plots of instantaneous LFP traces for lf and hf bursts (see (D)) for a TRN cell during baseline (top) and ArchT (bottom) conditions. Colors indicate the amplitude values of the standardized LFP traces. Rows represent individual burst events. 0 marks the first spike in the burst. White asterisks mark the peak values of the individual instantaneous LFP traces. Red arrow shows the time lag between the instantaneous LFP peak and the first spike for an example burst. For the better comparison, equal number of events for all firing categories and conditions were randomly selected. (H) Average LFP traces (spike triggered averages, STAs) for hf (blue) and lf (orange) bursts for the same cell as in (G) during baseline (top) and ArchT (bottom) conditions. STA peak for hf bursts is indicated with dashed line. STA amplitudes for hf bursts and lf burst at this value are labelled by red arrowheads. Note diminished difference between the STA of hf and lf bursts in the ArchT condition. (I) STA amplitudes for hf and lf TRN bursts at the STA peak of hf bursts at baseline (blue line) vs ArchT (blue dashed line) conditions for the same cell. (J) Same as (I) for the lag between the individual LFP peaks and the first spikes of the bursts. (K) Same as (H) for n=6 cells. upper panel: baseline; lower panel: ArchT; population averages ±SEM are indicated. (L) Same as (I) for n=6 TRN cells, STA amplitudes for hf vs. lf bursts at the STA peaks for hf bursts during baseline (upper right panel) vs ArchT (lower right panel) conditions. upper right panel: Box plot of the values in the upper left panel (Student’s paired sample t-test) lower right panel: Box plot of the values in the lower right panel (Student’s paired sample t-test). (M) Same as (J) for n=6 TRN cells. Average lag between the individual LFP peaks and the 1st spike of the bursts during baseline (upper left panel) vs ArchT (lower left panel) conditions. upper right panel: Box plot of the values in the upper left panel. (Student’s paired sample t-test) lower right panel: Box plot of the values in the lower right panel (Student’s paired sample t-test) au: arbitrary unit; ic: internal capsule

Upon ArchT activation, however, the significant correlation between the aIBF of the bursts and the magnitude of the instantaneous LFP slope observed during baseline activity disappeared in 6 out of 7 cells (Pearson correlation; baseline: R=0.21±0.03; P<0.01 for 5 cells, P<0.05 for 2 cells; optogenetic perturbation: R=0.08±0.02; P<0.05 for 1 cell, P>0.05 for 6 cells) (Fig7, E-F). The significant difference between STA peak amplitudes for high frequency (hf) and low frequency (lf) bursts observed at baseline activity got also diminished. (Student’s paired sample t-test; baseline: 0.66±0.17 au vs. −0.14±0.22 au p=0.047; optogenetic perturbation: 0.46±0.11 au vs. 0.22±0.12 au, p=0.1657) (Fig7, G-I, K-L). During baseline activity hf bursts occurred significantly earlier than lf bursts relative to the instantaneous LFP peaks (Student’s paired sample t-test; lag of 1^st^ spike if 0 is the LFP peak: −0.18±6.41 ms vs. 15.7±4.15 ms p=0.0056) (Fig7, G, J, M). This difference disappeared upon perturbation of L5 inputs (Student’s paired sample t-test; lag of 1^st^ spike if 0 is the LFP peak: 13.9 ±7.7 ms vs. 7.7±6.48 ms p=0.1389) (Fig7, G, J, M). During ArchT activation the wavelet and FFT power spectra of the frontal cortical LFPs (FigS4, E-F), or the average waveform of the fast LFP events did not change (FigS4, G) indicating that the observed effects were not due to an overall change in cortical activity. In control experiments (n=4 cells in n=2 mice from which n=1 from Thy1-ChR2-EYFP and n=1 from Thy1-Cre line) we did not observe alterations in the TRN spiking activity – cortical LFP correlation (FigS5, A-E).

These data demonstrate that while slight perturbation of L5 terminals with ArchT in the TRN did not have major effect on the basic firing properties of the anterior TRN cells, it clearly disrupted the instantaneous correlation between the ongoing cortical activity and TRN spiking, indicating an instrumental role of the L5 input to control the readout of the cortical activity by the TRN.

### L5-TRN pathway mediates both feedforward and lateral inhibition

What are the thalamic targets of the TRN cells conveying the integrated activity of L5 neurons? To study this, we reconstructed the complete axon arbour of neurobiotin-filled TRN cells optogenetically tagged via their L5 inputs (Fig1, J; Fig6, B; Fig7, B; FigS5, B) (n=18 cells in n=13 mice, from which n=7 Thy1-ChR2-EYP, n=6 Rbp4-Cre) (Fig8, A-F, FigS6). All TRN neurons targeted thalamic nuclei known to be connected with the frontal cortex (Jones, 2007). These include the ventral lateral nucleus (VL): n=8; ventral anterior nucleus (VA): n=1; (VM): ventral medial nucleus; intralaminar complex (IL): n=6; parafascicular nucleus (Pf): n=2; mediodorsal nucleus, lateral part (MDL): n=2; mediodorsal nucleus, central part (MDC): n=1; mediodorsal nucleus, medial part (MDM): n=1; submedius nucleus (Sub): n=3), (Fig8, C-D). Interestingly 9 out of the 18 cells had more than one target nuclei (VA-VM: n=1; VM-Sub: n=1; VM-VL-Sub: n=1; VL-VM-IL: n=1; VM-IL: n=1; MD-IL: n=3; VL-AV: n=1). Although there was a loose topography regarding the position of the cell bodies and the axonal targets, TRN neurons with different targets could be found intermingled (Fig8, C; FigS7, A-B).

**Figure 8.**
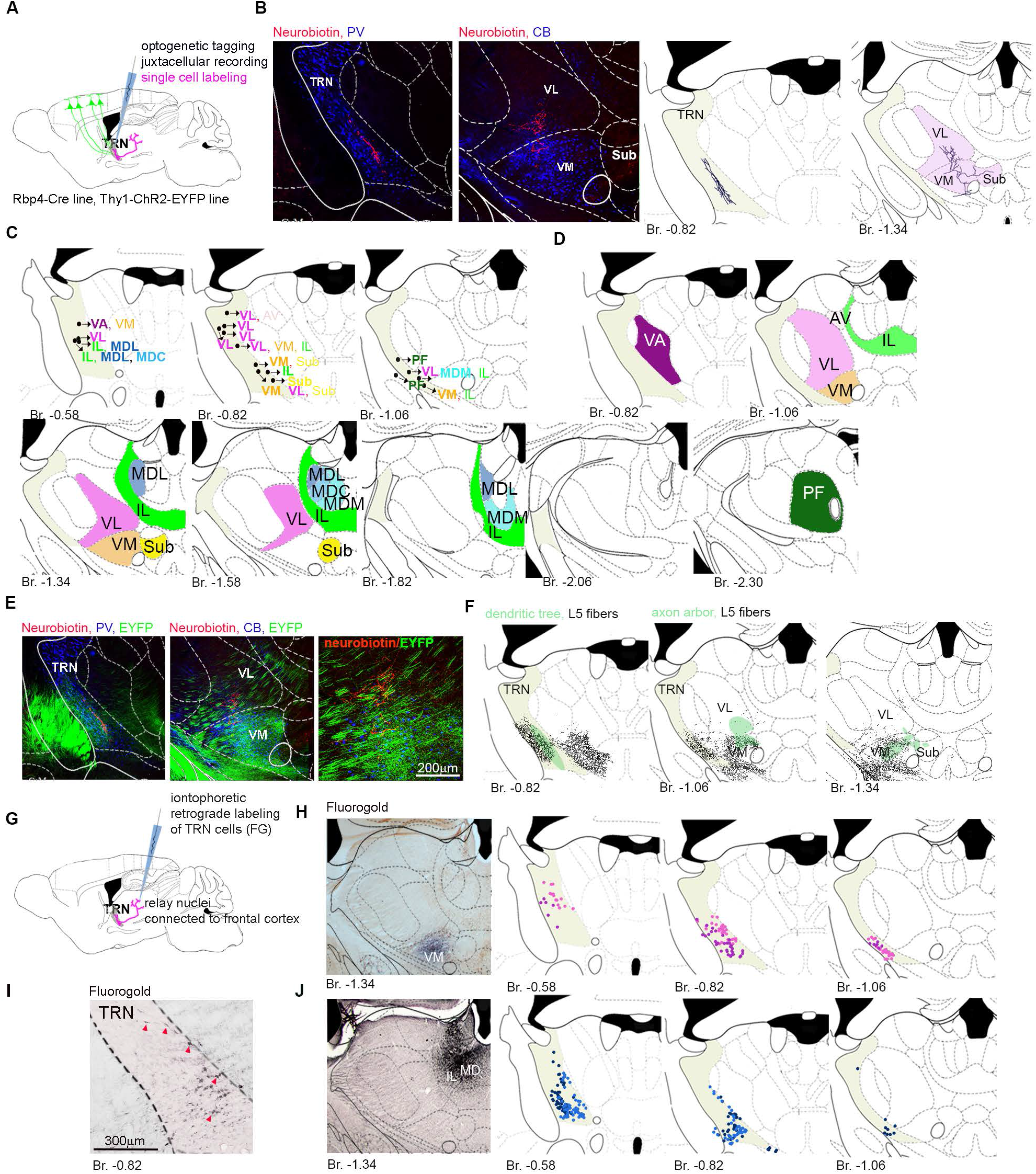
TRN cells receiving L5 input target relay nuclei connected to the frontal cortical areas. (A) Schematics of experimental design to label the output of optogenetically tagged and juxtacellularly recorded TRN neurons. (B) An example of a neurobiotin (red) filled TRN neuron. Left panel: confocal image of the cell body and the dendrites (red) in the TRN (PV+, blue). Middle left panel: confocal image of the axon arbor (red) in the VM (CB+, blue) and VL (CB-) relay nuclei. Middle right panel: Reconstructed soma and dendritic tree in the TRN. Right panel: Reconstructed axon arbor in the relay nuclei (target nuclei are labelled with pale red). TRN: thalamic reticular nucleus VL: ventral lateral nucleus; VM: ventral medial nucleus; Sub: submedius nucleus; Br: Bregma; TRN is labelled in grey. (C) Cell body position of reconstructed TRN neurons (n=18 in n=14 mice, from which n=7 Thy1-ChR2-EYP, n=7 Rbp4-Cre) driven from the frontal cortex. Arborization zone of the labelled neurons are indicated by the abbreviations of the target nuclei. Font colors match the colors of the target nuclei on (D). Primary target nuclei are labelled in bold. VA: ventral anterior nucleus; VL: ventral lateral nucleus; VM: ventral medial nucleus; AV: anteroventral nucleus; IL: intralaminar complex; Pf: parafascicular nucleus; MDL: mediodorsal nucleus, lateral part; MDC: mediodorsal nucleus, central part; MDM: mediodorsal nucleus, medial part; Sub: submedius nucleus; Br: Bregma; TRN is labelled in grey. (D) Target nuclei of the TRN neurons shown in (C). For the extent of individual axon arbors see Figure S6. TRN is labelled in grey. (E) Left panel: an example of optogenetically tagged and neurobiotin filled TRN neurons from an Rbp4-Cre mice surrounded by virus labelled L5 fibers originating from the M1/M2 cortex. Middle panel: Axon arbor of the same neuron together with the L5 fibers originating from the M1/M2. Right panel: Higher magnification for the middle panel. Axon arbor of the labelled TRN cell and the L5 afferents overlap in the VM, however, VL contains only TRN but no L5 terminals indicating an open loop condition. (F) Schematics depiction of (E). Black, L5 collaterals; green, dendritic and axon arbour of the TRN neuron. For more examples see Figure S8. (G) Experimental design. (H) Left panel: Injection site in the VM. Middle and right panels: Position of retrogradely labelled somata in the TRN. n=2 mice, labelled with different shades. TRN is labelled with grey. (I) Retrogradely labelled cells in the anterior TRN after retrograde tracer injection to the MD. Red arrowheads indicate some of the labelled cells. (J) Left panel: Injection site in the MD. Middle and right panels: Position of retrogradely labelled somata in the TRN. n=2 mice, labelled with different shades. TRN is labelled with grey. AV: anteroventral nucleus; Br: Bregma; FG: fluorogold; IL: intralaminar coplex; MD: mediodorsal nucleus; MDC: mediodorsal nucleus, central part; MDL: mediodorsal nucleus, lateral part; MDM: mediodorsal nucleus, medial part; Pf: parafascicular nucleus; Sub: submedius nucleus; TRN: thalamic reticular nucleus; VA: ventral anterior nucleus; VL: ventral lateral nucleus; VM: ventral medial nucleus

In accordance with these data, retrograde labelling experiments (n=4 mice) confirmed that TRN cells targeting frontal cortex related relay nuclei (VM, MD) are in the anterior, L5 recipient part of TRN (Fig8, G-J).

To resolve whether the TRN transmits feed forward or lateral inhibition we reconstructed the axon arbors of TRN cells in the thalamus together with the L5 fibers via which they were activated in the Rbp4-Cre mice (n=5 cells in 4 animals). We examined whether TRN axons are inside (indicating feed forward inhibition) or outside (lateral inhibition) the labeled cortical terminal field in the thalamus (Fig8, E-F; FigS7, E-H).

In three cases large proportion of the TRN axon arbor was outside the termination zone of the cortical L5 fibers, implying that lateral inhibition can be significant in the L5-TRN pathway. In all three cases the L5 axons were confined to VM whereas TRN axons innervated VL. In the other two cases (VM and Pf nuclei) TRN axonal targets were completely within the L5 zone (n=2 cells), indicating feed forward inhibition in case of these neurons.

In one of the *in vivo* electrophysiology experiments, two TRN cells could be recorded and filled simultaneously in the anterior TRN (FigS7, A). The cell bodies were in close vicinity of each other (within 100 µm), but the two TRN cells targeted two different relay nuclei (MD and VM respectively) (FigS7, B). Upon fast cortical events, the firing of the two cells got tightly correlated (FigS7, C). STA for the spikes which were paired with a spike of the other cell within a 5 ms time window had a higher amplitude and narrower peak compared to the STA of the spikes which were less synchronous with the activity of the other cell (FigS7, D).

These data show that L5 neurons in one cortical location can have widespread inhibitory action in the thalamus via their connection with TRN and suggests that synchronous cortical events can synchronize the activity of multiple relay nuclei via the L5-TRN output.

## Discussion

In this paper we described and characterized a specific and topographically organized pathway originating from the frontal cortical L5 PT cells, which selectively targets the anterior TRN. The data showed that via integrating multiple L5 inputs the exact spike output of the TRN cells provided a sensitive measure of synchronous cortical activity. The output of frontal L5-driven TRN activity reached thalamic regions connected to the frontal cortices. These data indicate that cortical control of thalamic activity is region-specific, and that frontal L5-TRN projection can be instrumental in sculpting thalamic activity in widespread frontal cortical functions involving synchronous cortical firing.

Until recently, the organization of corticothalamic connections was considered canonical (Harris and Shepherd, 2015; Sherman and Guillery, 2013; Thomson, 2010). All cortical regions were reported to send both L6 and L5 projections to the thalamus but only L6 corticothalamic axons have been shown to establish synaptic connections in the TRN. Indirect evidence from previous reports indicated the presence of L5 synaptic input in the TRN (Prasad et al., 2020; Rockland, 1998; Zikopoulos and Barbas, 2006). However, none of the these works provided direct, conclusive evidence for a monosynaptic L5-TRN connection, cell type specificity or regional variability of its source (Guo et al., 2018) and their physiological features. Here we clearly demonstrate monosynaptic connection selectively from L5 PT cells (Figure 2) of the frontal cortex to the TRN using both morphological and physiological methods, and so we provide direct evidence for qualitative differences between cortical regions regarding the way they recruit intrathalamic inhibition (Fig1).

We demonstrated that along the specific L5-TRN projection from the frontal cortex the canonical L6-TRN input forms a highly convergent, parallel pathway in the anterior TRN (Fig3, A-C). The two cortico-TRN pathways display divergent morphological and functional properties (Fig3-4). The difference in the organization of L6-TRN and L5-TRN pathways is comparable to those of the L6-thalamic and L5-thalamic pathways (Alexander et al., 2006; Crandall et al., 2015; Ohara, 1988; Sherman and Guillery, 2013; Wilson et al., 1984) confirming complementary roles of the two projections. The larger volume of L5-TRN boutons can be largely attributed to the presence of multiple mitochondria suggesting intensive synaptic activity (Vos et al., 2010). The diameter of the dendrites negatively correlates with the distance from the soma. Thus, the larger diameter of the postsynaptic dendrites in case of the L5-TRN synapses suggests more proximal and more effective synaptic connection. The presence of spine synapses, the complex PSD morphology (Fig3) and the elevated NMDA/AMPA ratio (Fig4) may indicate a higher potential for synaptic plasticity at the L5 to TRN synapses (Astori and Lüthi, 2013; Hering and Sheng, 2001).

Our data shows that individual TRN cells effectively integrate the activity of multiple presynaptic L5 cells. While individual L5 EPSCs were small *in vitro* (Fig4) optogenetically activated L5 neurons could reliably fire TRN cells *in vivo* (Fig1). The response probability and the exact spike output of the optogenetically evoked TRN responses *in vivo* correlated with the size of recruited L5 population activity (Fig6, FigS1). Moreover, during spontaneous activity the firing pattern of 61.3 % of the anterior TRN neurons provided a gradual, L5-mediated read-out of the magnitude of synchronous cortical activity which could be detected in the LFP recordings as fast, high amplitude transients (Fig6). While single spikes were uncorrelated with the cortical LFP, bursts were specifically coupled to the fast cortical transients, and the burst properties of the TRN neurons significantly correlated with the magnitude of the instantaneous LFP slope (Fig6). Bursts have been described both in awake and sleep states in the anterior TRN (Marlinski and Beloozerova, 2014). Although bursts are often viewed as stereotypical all-or-none events, experimental evidence shows considerable variety in the TRN burst properties, which can correlate e.g. with the complexity of the behavior (Marlinski and Beloozerova, 2014) or with the parameters of the corticothalamic oscillations (Barthó et al., 2014). Furthermore, Kepecs and colleagues showed that bursts tend to occur at the positive slope of the synaptic input signals and that burst properties can code the magnitude of the signal slope (Kepecs et al., 2002). Based on this, we propose that the exact spike output and burst pattern of TRN neurons will code the level of synchronous L5 activity in the cortex.

Optogenetic perturbation of L5 fibers in the TRN further confirmed the critical role of L5-TRN input to transmit fast changes in cortical activity to anterior TRN cells (Fig7). Upon ArchT-mediated disruption of L5-TRN inputs, the correlation between TRN burst properties and the instantaneous cortical LFP-activity was disturbed. Earlier work showed that ArchT activation in presynaptic terminals does not result in a clear inhibition of synaptic transmission, but rather in a mixture of decreased probability of AP evoked release and increased probability of spontaneous synaptic release (Mahn et al., 2016). In line with this we did not observe long term decrease in firing rates of the TRN neurons upon sustained ArchT activation of their L5 inputs. Decoupling of presynaptic activity and precise transmitter release in the L5 terminals via ArchT, however, was sufficient to disrupt the correlation between the cortical and the TRN-activity. These data show that precise and effective integration of L5 output is required to convert cortical activity to a TRN AP output pattern.

What might be the significance of the exact TRN spike output for the postsynaptic thalamocortical neurons? TRN bursts can increase the IPSC magnitude in the postsynaptic relay cells compared to single TRN APs (Cox et al., 1997). Mechanism of burst IPSCs is unlike that of single APs since GABA released during bursts can recruit nonsynaptic GABA-A receptors, which results in a significantly different inhibitory charge and kinetics (Herd et al., 2013; Rovo et al., 2014). In freely sleeping conditions, the exact number of spikes/TRN burst changes stereotypically during sleep spindles and cycle-by-cycle reduction in spike/burst was suggested to be a major determinant of terminating this sleep transient (Barthó et al., 2014). Thus, the impact of different TRN spike patterns on relay cell firing and signal integration is clearly significant but certainly needs further investigation.

Synchronous activity can arise locally or from multiple regions of the frontal cortex. In our viral tracing experiments, L5 axons from the neighbouring frontal cortical territories showed clear segregation (Fig5). Our data was in a good agreement with the single cell reconstruction data from the Mouse Light Neuron Browser and with the paper of Lozsadi (Lozsadi, 1994) which reported loose but clear topography in the cingulate cortex - TRN pathway. Topographical termination of frontal L5 fibres in the TRN suggested that TRN cells at a given spatial position may integrate inputs from a relatively narrow cortical territory. The extensive dendritic tree of the L5-driven TRN neurons (Fig1, Fig6, Fig7, Fig 8) however may extend across multiple termination zones so TRN cells could integrate more global synchronous cortical activity. Indeed, our experiments demonstrated that while TRN cells showed preferential activation from specific frontal cortical areas, they could be activated from multiple frontal cortical territories (Fig5).

Under our experimental conditions firing properties of 38.7% of the recorded anterior TRN neurons did not show significant correlation with the frontal LFP activity. These neurons had also significantly lower firing and burst rates and were spatially mixed with the neurons which showed significant correlation with the cortical LFP (FigS4). Recent investigations (Clemente-Perez et al., 2017; Li et al., 2020; Martinez-Garcia et al., 2020) demonstrated the presence of at least two morphologically and physiologically different cell types in the TRN. Whether the two anterior TRN populations in our experiments corresponds with these previously described groups requires further investigation.

Tracking the axons of L5 driven TRN cells clearly showed that, via their TRN collaterals, L5 neurons of the frontal cortex can have widespread inhibitory action in large thalamic regions related to the frontal cortex (Fig8). Our experiments revealed anatomical basis for feed forward inhibition in the VM and Pf nuclei. In contrast, several L5-recipient TRN cells innervated the VL nucleus, which, as a first order nucleus, does not receive L5 input from the frontal cortex (Shepherd and Yamawaki, 2021) (Fig8, FigS6) indicating lateral inhibition and crossmodal interactions in the case of VL. L5 driven TRN cells frequently innervated multiple thalamic nuclei, a rare feature of TRN cells (Pinault and Deschenes, 1998), and TRN cells targeting different thalamic nuclei displayed correlated activity during fast cortical transients (FigS7). This suggests that synchronous cortical events can synchronize the activity of multiple relay nuclei via the L5-TRN output.

Feed forward (Buzsàki and Eidelberg, 1981) and lateral inhibition are fundamental mechanisms of the neuronal circuits which, among other things, are pivotal for gain control (Halassa and Acsády, 2016), synchronization of high frequency activity (Zemankovics et al., 2013), frequency dependent signal transfer (Morl et al., 2004) or receptive field tuning (Swadlow, 2002). Our data suggest that corticothalamic L5 circuits are heterogeneous in this respect. We show here that in contrast to sensory corticothalamic information transfer the vast array of frontal cortical functions utilize an additional, powerful form GABAergic mechanism at the level of thalamus. Since frontal cortex is implicated in diverse neurological conditions (e.g. Parkinson’s disease, epilepsy, chronic pain) and thalamic neurons respond robustly to TRN inhibition, frontal L5-TRN projection characterised here may potentially play critical role in establishing and/or maintaining these pathological conditions.

## Ackowledgements

We thank the excellent technical assistance of Krisztina Faddi and Győző Goda. The authors thank the Light Microscopy Center and the Virus Technology Unit of the IEM to provide microscopy and technical support and Balázs Hangya and László Bíró for comments and discussions on the manuscripts. We are grateful for Ofer Yizhar for kindly providing us the AAV-DFO-ChR2-eYFP virus, for Balázs Rózsa for the Thy1-Cre animals and for Hajnalka Bokor for performing the VM experiments. This work was supported by the ERC (FRONTHAL, 742595 to L.A.).

## Author contribution

GV, AL participated in the in vitro investigations and performed review editing. EB and BT perfomed morphological and EB also in vivo investigations. NH participated in data curation, formal analysis, programming, morphological and in vivo investigations. NH and LA formulated the research idea, supervised the research, developed the methodology and visualization and wrote the original manuscript, LA was responsible for funding acquisition and validation.

## Star methods

### RESOURCE AVAILABILITY

#### Lead contact

Further information and requests for resources and reagents should be directed to and will be fulfilled by the lead contact, László Acsády.

#### Materials availability

This study did not generate new unique reagents.

#### Data and code availability

The datasets and the scripts used in study are available from the Lead Contact upon reasonable request.

### EXPERIMANTAL MODEL AND SUBJECT DETAILS

#### Animals

All animal use was approved by the Animal Welfare Committee of the Institute of Experimental Medicine, Budapest, in accordance with the regulations of the European Community’s Council Directive of November 24, 1986 (86/609/EEC). The experiments were approved by the National Animal Research Authorities of Hungary (PE/EA/877-7/2020). Mice were maintained on a 12 hr light/dark cycle, and food and water was provided ad libitum. All mice were healthy with no obvious behavioral phenotypes. For all mouse studies, adult male mice were used, and mice were randomly allocated to experimental groups. C57Bl/6J -Tg (Rbp4-Cre) (MGI:4367067, STOCK Tg(Rbp4-cre)KL100Gsat/Mmucd, ID 031125-UCD) (Gong et al., 2007), C57Bl/6J-Tg (Thy1-ChR2-YFP) (JAX stock #007612) (Arenkiel et al., 2013), FVB/AntFx-Tg (Thy1-Cre) (JAX stock #006143)(Dewachter et al., 2002) and Bl6Fx –Tg (Ntsr1-Cre) (MGI:3836636) (Gong et al., 2007) mice were mice were obtained from The Jackson Laboratory.

### METHODS DETAILS

#### Surgery

Mice were anesthetized with an intraperitoneal injection of ketamine-xylazine (ketamine, 83 mg/kg; xylazine, 3.3 mg/kg body weight) and placed inside a stereotactic apparatus. Sleep depth was monitored throughout the surgery, and additional dose (ketamine, 28 mg/kg; xylazine, 1.1 mg/kg body weight) of anaesthetic was applied intramuscularly if necessary.

##### Viral injections

Virus injections for anatomical and optogenetic experiments were performed on adult Rbp4-Cre, Thy1-Cre and Ntsr1-Cre mice. AAV5.EF1a.DIO.hChR2(H134R)-eYFP.WPRE.hGH (based on Addgene plasmid #20298, UNC Vector Core) AAV5.EF1.dflox.hChR2(H134R)-mCherry.WPRE.hGH (based on Addgene plasmid #20297, UNC Vector Core) AAV5.CAG.Flex.ArchT-GFP (based on Addgene plasmid # 28307, UNC Vector Core), AAV.DFO.ChR2-eYFP (Barsy et al., 2020) viruses were injected in the right side neocortex or brainstem (200 nl, 1 nl/sec) using pipettes pulled from borosilicate glass capillaries. The stereotaxic coordinates were the following (AP and ML taken from the bregma, DV taken from the brain surface).

Cortical injections resulting in L5 collaterals in the TRN: M2: AP +2 mm, ML +0.5 mm, DV – 0.7 mm (in case of double injections to M2, M2 anterior: AP: 2.5 mm, ML 0.7 mm, DV - 0.7 mm; M2 posterior: AP +1.5 mm, ML +0.7 mm, DV: −0.7 mm); M1: AP +2mm, ML + 2mm, DV −0.7mm; ALM: AP + 2.7 mm, ML + 1.5 mm, DV - 0.7mm; LO/VO: AP +2.2, ML +1.2 mm, DV −0.7 mm; PrL: AP +2 mm, ML +0.4 mm, DV −1.2 mm; Cg/RS: AP −1 mm, ML +0.5 mm, DV +0.7 mm or AP −2mm, ML −0.5 mm, DV −0.7 mm;

Cortical injections resulting in no L5 collaterals in the TRN: S1J: AP +1.1 mm, ML +2.6 mm, DV −1; S1BF: AP −1.3 mm, ML +3 mm; DV: −0.7 mm; S2: AP −0.1 mm, ML +3.5, mm DV −1.7 mm; GI/DI: AP: +0.1 mm, L +3.6 mm, DV −2.3 mm;

Brainstem: AP −4.4 mm, ML 0.8 mm, DV −4.2 mm.

##### Tracer injections

Retrograde tracer injections were performed on adult wild type littermates of Rbp4-Cre mice. Fluorogold (FG) was injected iontophoretically (0.5 μA; 2 s on/off period; 10 min duration), in the right side of the thalamus using pipettes pulled from borosilicate glass capillaries. The stereotaxic coordinates were the following (AP and ML taken from the bregma, DV taken from the brain surface). VM: AP −1.3 mm, ML 0.8 mm, DV −4.2 mm; MD AP −1.3 mm, ML 0.5 mm, DV −2.9.

#### Histology

Mice were perfused with a fixative containing 4% paraformaldehyde (PF) and 0.1% glutaraldehyde in PB (0.1 M phosphate buffer). After perfusion, brains were removed from the skull, and coronal sections (50 μm thick) containing the cortex, thalamus or brainstem were cut with vibratome. To permeabilize the cell membranes sections were incubated in sucrose (30%) overnight, followed by freeze-thawing over liquid nitrogen. Fluorogold was visualized with a rabbit anti-Fluorogold antibody (1:10,000, Chemicon) followed by biotinylated goat anti-rabbit (1:300 Vector Laboratories) and avidin biotinylated horseradish peroxidase complex (ABC, 1:300 Vector Laboratories). Nickel-intensified 3,3′-diaminobenzidine (DABNi, bluish-black reaction product) was then used as a chromogen. EYFP and mCherry fluorescent labels were intensified via chicken anti-eGFP antibody (1:5000, Life Technologies) followed by goat anti-chicken-Alexa488 antibody (1:500, Molecular Probes) and rabbit anti-mCherry antibody (1:3000, BioVision) followed by donkey anti-rabbit-Cy3 antibody (1:500, Jackson) respectively. Neurobiotin content of juxtacellularly labelled cortical, TRN and VM cells was visualized by Cy3-Streptavidin (1:500, Jackson). For reconstructing axonal and dendritic arbours of TRN cells, DABNi staining was developed in the neurobiotin labelled cells after applying avidin biotinylated horseradish peroxidase complex (ABC, 1:300 Vector Laboratories, see above). Cortical synaptic terminals were labelled by Vesicular glutamate transporter 1 staining (rabbit anti-Vglut1 antibody, 1:10 000, Millipore; donkey anti-rabbit-Cy3 antibody, 1:500, Jackson). Boundaries of TRN and higher order relay nuclei were determined by parvalbumin (PV, mouse anti-PV antibody, 1:2000, Sigma; donkey anti-mouse-Cy5, 1:500, Jackson) and calbindin (CB, rabbit anti-CB antibody, 1:2000,; donkey anti-rabbit-Cy5, 1:500, Jackson) staining respectively. The results of the light microscopic experiments were obtained with either a Zeiss Axionplan 2 fluorescent microscope and photographed by a digital camera (Olympus DP70), or with a Zeiss Axio Imager M1 microscope coupled to an AxioCam HrC digital camera, or with a Nikon AR1 confocal microscope.

#### Electron microscopy

M1/M2 cortices of adult Rbp4-Cre or Ntsr1-Cre mice were injected with AAV-EF1a-DIO-ChR2-EYFP virus. Two weeks post-surgery, mice were perfused with a fixative containing 4% paraformaldehyde (PF) and 0.1% glutaraldehyde in PB (0.1 M phosphate buffer). After perfusion, brains were removed from the skull, and coronal sections (50 μm thick) containing the thalamus were cut with vibratome. To permeabilize the cell membranes sections were incubated in sucrose (30%) overnight, followed by freeze-thawing over liquid nitrogen. Labeled fibers were visualized with a nickel-intensified 3,3′-diaminobenzidine (DABNi) reaction resulting in a bluish-black reaction product. Sections were first incubated with rabbit anti-eGFP antibody (1:2000, Life Technologies) overnight, followed by biotinylated b-SP donkey anti-rabbit antibody (1:300; Jackson). Sections were incubated with avidin biotinylated horseradish peroxidase complex (ABC; 1:300; Vector Laboratories) for 2 h and finally, the staining was developed with DABNi. Sections were treated with OsO4, dehydrated in ethanol and propylene oxide, and embedded in Durcupan. During dehydration, the sections were treated with 1% uranyl acetate in 70% ethanol. Selected blocks containing labeled L5 terminals in the TRN were reembedded, 60-nm-thick (silver color) ultrathin sections were cut with an Ultramicrotome, and sections were mounted on copper grids. Serial electron micrographs were taken with a Megaview digital camera running on a HITACHI 7100 electron microscope. Reconstruct™ software was utilized to reconstruct the terminals in 3D. The sections were aligned with the ‘linear alignment’ function that can partially compensate for individual section distortions caused by sectioning. Cell membranes of pre- and postsynaptic structures, postsynaptic densities (PSD) and mitochondria in the synaptic boutons were reconstructed. For further details on reconstructions see. For 2D measurements, Fiji software was used. The minor diameters of dendrites targeted by eGFP-positive boutons were measured in 3 non-consecutive sections and averaged. Bouton volume, PSD area and number of mitochondria were calculated in boutons where consecutive sections containing the full extent of the given structure were preserved, and the ultrastructure (Wanaverbecq et al., 2008) of the tissue was appropriate.

#### *In vivo* electrophysiology

##### Anesthesia and surgery

Adult male Thy1-ChR2-EYFP, Rbp4-Cre and Thy1-Cre mice (20-30g) were used for the experiments. In the case of Rbp4-Cre and Thy1-Cre mice, electrophysiology experiments were carried out 2-4 weeks after the virus (AAV5.EF1a.DIO.hChR2(H134R)-eYFP.WPRE.hGH or AAV5.CAG.Flex.ArchT-GFP) injection to M2 and M1(200-200nl). Surgeries and experiments were done under ketamine/xylazine anesthesia. Initially, mice received intraperitoneal injection of ketamine (50 mg/kg) and xylazine (4 mg/kg). For n=3 mice we used the doses used at surgeries for virus or tracer injections (ketamine, 83 mg/kg; xylazine, 3.3 mg/kg body weight). This higher dose however, induced a deep anaesthesia characterized by a frontal cortical LFP dominated by a regular, slow (1-4 Hz) component, and by a predominantly tonic firing of TRN neurons. For this reason, mice receiving this higher dose of ketamine/xylazine were excluded from the analyses examining cortical LFP-TRN firing correlation. For optogenetic perturbation experiments using ArchT, and for their control experiments 36 mg/kg ketamine and 2.9 mg/kg xylazine was used. For the maintenance of the anesthesia, intramuscular injection of ketamine/xylazine (1/3 of the initial amount) was given every 30-50 min during the entire duration of the experiments.

##### In vivo juxtacellular recording and labeling, local field potential (LFP) recording

Bipolar LFP electrodes (FHC, resistance ∼1 MΩ) were placed into the frontal cortex of mice (AP: +2.5, ML: 1 mm from Bregma). The recorded signal was amplified, band-pass filtered from 0.16 Hz to 5 kHz (Supertech) and digitized at 20 kHz (micro 1401 mkii, CED). Cortical L5, TRN or VM single unit activity was recorded by glass microelectrodes (*in vivo* impedance of 20-40 MΩ) pulled from borosilicate glass capillaries (1.5 mm outer diameter, 0.75 or 0.86 inner diameter, Sutter Instrument Co) and filled with 0.5 M K^+^-acetate and 2% neurobiotin (Vector Laboratories). Electrodes were lowered by a micromanipulator (Scientifica) to the target area (Cortex: AP +2 mm, ML +0.5 mm, DV −0.5-0.9 mm; TRN: AP −0.7 mm, ML 1.6 mm, DV −2.3-4 mm, VM: AP −1.3 mm, ML +0.9 mm, DV −3.8-4.3 mm; AP and ML taken from the bregma, DV taken from the brain surface). Neuronal signals were amplified by a DC amplifier (Axoclamp 2B, Axon Instruments/Molecular Devices), further amplified and filtered between 0.16 Hz and 5 kHz by a signal conditioner (LinearAmp, Supertech). Neuronal signals were recorded by Spike2 7.0 (CED). Juxtacellular labeling of the recorded neurons was done as described previously (Pinault, 1996). For histological analysis see Method details/ Histology.

#### *In vivo* optogenetics

For optogenetic activation of L5 cells Adult male Thy1-ChR2-EYFP and Rbp4-Cre mice were used. In the case of Rbp4-Cre mice, AAV5.EF1a.DIO.hChR2(H134R)-eYFP.WPRE.hGH virus was injected (See Method Details/Surgery). Juxtacellular recording and cortical LFP recording was carried out as in Method Details/*In vivo* electrophysiology. The skull was thinned above the M2/M1 area of the right cortical hemisphere, where the optic fiber (100 µm, 0.22 NA) was positioned (M2: AP +2 mm, ML + 0.5 mm; M1: AP +2mm, ML + 2mm, from the bregma). The laser beam was generated by a 473 nm DPSS laser (Laserglow Technologies). Laser power at the tip of the optic fiber by different switch positions was measured both before and after each experiment with a photometer (Thorlabs). In addition, laser power was monitored throughout the experiment via a photometer (Thorlabs) built in the laser path. TRN cells targeted by labelled L5 fibers were find via optogenetic tagging by applying test laser pulses (5 ms, 10mW). After recording the baseline activity for 300 s, 5 stimulus trains of 10×1Hz 1ms (in case of two Thy1-ChR2-EYFP animals) or 5ms (in all other cases) long pulses were applied which were generated by Spike2 7.0 software (CED).

For optogenetic perturbation of L5 to TRN inputs adult male Thy1-Cre and Rbp4-Cre mice were injected with AAV5.CAG.Flex.ArchT-GFP virus (See Method Details/Surgery). For control experiments Thy1-ChR2-EYFP and Thy1-Cre mice were used. Juxtacellular recording and cortical LFP recording was carried out as described in Method Details/*In vivo* electrophysiology. Custom made mirror tip optic fibers (200 µm, 0.37 NA, Doric) were lowered to the target area in the TRN (AP −0.5 mm, ML 0.8 mm, DV −4 mm; AP and ML taken from the bregma, DV taken from the brain surface) in 20° angle, with the mirror facing towards the TRN, preventing the light to spread to the neighbouring relay nuclei. The laser beam was generated by a 589-594 nm DPSS laser (Laserglow Technologies). Laser power at the tip of the optic fiber was measured both before and after each experiment with a photometer (Thorlabs). TRN cells targeted by labelled L5 fibers were found via optogenetic tagging by applying test laser pulses (5s, 10 mW). A slight and transient drop in in the firing frequency of the targeted cells could be observed. In case of the control experiments cells near the optic fiber tip were found by optogenetic tagging with 5ms, 5 mW test pulses (473 nm, Thy1-ChR2-EYFP mouse) or post hoc by measuring the distance between the position of the soma and the optic fiber tip (<200 µm, Thy1-Cre mouse). After recording the baseline activity for 300 s, 60 cycles of 5s long 589-594 nm laser pulses (laser ON) followed by 10 s laser OFF period (10s) were applied. Position of the optic fiber, labelled L5 terminals and TRN somas were verified for each experiment after histological processing of the brains (see Method Details/Surgery).

#### *In vitro* electrophysiology

##### Brain slicepreparation

Rbp4-Cre and Ntsr1-Cre mice were sacrificed 3 to 4 weeks post viral injection. Briefly, mice were anesthetized with isoflurane and their brains quickly extracted. Acute 300-µm-thick sections were sliced using a sliding vibratome (Histocom, Zug, Switzerland) while submerged in ice-cold oxygenated sucrose solution (which contained in mM: 66 NaCl, 2.5 KCl, 1.25 NaH_2_PO_4_, 26 NaHCO_3_, 105 D(+)-saccharose, 27 D(+)-glucose, 1.7 L(+)-ascorbic acid, 0.5 CaCl_2_ and 7 MgCl_2_). Slices were then stored in a recovery solution (in mM: 131 NaCl, 2.5 KCl, 1.25 NaH_2_PO_4_, 26 NaHCO_3_, 20 D(+)-glucose, 1.7 L(+)-ascorbic acid, 2 CaCl_2_, 1.2 MgCl_2_, 3 myo-inositol, 2 pyruvate) at 35°C for 30 min and then at room temperature for 30 min before recording.

##### Whole-cell patch-clamp recording and optogenetic stimulation

The recording extracellular solution (containing in mM: 131 NaCl, 2.5 KCl, 1.25 NaH_2_PO_4_, 26 NaHCO_3_, 20 D(+)-glucose, 1.7 L(+)-ascorbic acid, 2 CaCl_2_, 1.2 MgCl_2_) was supplemented with 0.1 picrotoxin, 0.01 glycine when appropriate (Fig4, E), maintained at room temperature, constantly oxygenated and perfused in the recording chamber. Borosilicate glass pipettes (TW150F-4, World Precision Instruments) were filled with an intracellular solution containing (in mM): 140 KGluconate, 10 HEPES, 10 KCl, 0.1 EGTA, 10 phosphocreatine, 4 Mg-ATP, 0.4 Na-GTP, pH 7.3, 290 – 305 mOsm, supplemented with ∼2 mg/ml of neurobiotin (Vector Labs, Servion, Switzerland)) and showed a pipette resistance ranging from 2.5 to 5 MΩ for most recordings. NMDAR-mediated currents (Fig4, E) were recorded using an intracellular solution containing (in mM): 127 CsGluconate, 10 HEPES, 2 CsBAPTA, 6 MgCl_2_, 10 phosphocreatine, 2 Mg-ATP, 0.4 Na-GTP, 2 QX314-Cl, supplemented with ∼2 mg/ml of neurobiotin, pH 7.3, 290–305 mOsm.

Whole-cell patch-clamp recordings were performed as previously described (Vantomme et al. 2020). Briefly, passive cellular properties were measured immediately after graining whole cell access, while holding the TRN cell at −60 mV. L5 and L6 afferents were activated using whole-field blue LED (Cairn Res, Faversham, UK) stimulation (455 nm, duration: 0.1 to 1 ms, maximal light intensity 3.5 mW, 0.16 mW/mm2) during voltage-clamp recordings. The properties of the EPSCs were measured at −60 mV. Paired pulse stimulation at 1, 2, 5, 10 and 20 Hz were used to measure the short-term properties of L5 and L6-TRN synapses. The AMPA receptor mediated component was measured at −60 mV and blocked using 40 µM DNQX. Once fully blocked, the cell was clamped at +40 mV to record the NMDA receptor mediated component. D,L-APV (100 μM) was then use to fully block the evoked response.

### QUANTIFICATION AND METHOD ANALYSIS

Analysis of the *in vitro* electrophysiology data was performed by built-in and custom written Matlab code (Mathworks).

#### Spiking data

Spike detection was carried out by Spike2 7.0 software. Spiking data was downsampled to 10 000 Hz. We defined a burst as an action potential package with interspike intervals (ISIs) below 10 ms. Avarage intraburst frequency of a burst (aIBF) was calculated as the average of the reciprocal of all ISIs within the burst. Responses of TRN cells to L5 stimulation were defined as single spikes within 30 ms or bursts with the first spikes within 30 ms following the start of the laser pulse.

#### Cortical LFP data

LFP data was downsampled to 10 000 Hz, except for the fast Fourier transform (FFT) analysis, where cortical LFP was downsampled to 500 Hz. For the FFT analysis Welch’s power spectral density estimate was used. LFP traces were Z-scored. Wavelet calculations were performed using Morlet wavelet transform.

#### Spike triggered average (STA) analysis

STAs were calculated as the averages of frontal cortical LFP traces ± 100 ms around the TRN single spikes or around the first spikes of the TRN bursts, except for Supplementary Fig. 7, where all spikes were treated individually. TRN spiking events were selected into five categories. Single spikes and 4 burst categories according to the aIBFs. Boundaries of the four categories were defined by the 25th, 50th and 75th percentiles. STA peak amplitude was calculated first for the 4th burst category (high frequency-hf - bursts). Because STA peaks for the 1st burst category (low frequency –lf-bursts) were often less well defined, we calculated the STA amplitude for lf bursts at the position of the hf STA peak. For calculating the average lags of the LFP peaks from t=0 ms, peak amplitudes were calculated for individual LFP events, and their lags from 0 were averaged.

#### Correlation between cortical LFP slope and TRN burst properties

Instantaneous LFP slope for each burst event was calculated as the steepness of the linear fit to a 30ms section of the LFP data around the first spikes of the bursts. TRN neurons fired at different phases of the LFP sharp events. To compensate for this, and measure the steepness of the positive slope of the instantaneous LFP events, we choose the 30 ms LFP sections for the linear fit as the following:

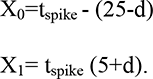

Where X_0_ is the start and X_1_ is the end of the selected LFP section in ms. t_spike_ is the time point for the first spike of the burst in ms, d is the lag of the STA (LFP average for all bursts) peak in ms.

Next, we defined the Pearson correlation coefficient (R) and the p values for the correlation between the aIBF value of the bursts and the slope of the instantaneous cortical LFP slope. To be able to compare p values, for the optogenetic perturbation experiments equal number of bursts were selected from the baseline and the laser recordings. Bursts were selected uniformly at random without replacement, and R and p values were calculated. This was repeated 200 times, and the averages of R and p values were calculated.

#### Sharp event detection

30 ms long sections of the (120-300 s long) LFP recordings were randomly selected 200 000 times, and the steepness of the linear fits for these sections were calculated. The steepest LFP sections with steepness values larger than the 98^th^ percentile were selected. Next, we searched for the peaks of these fast LFP events. Maximum amplitudes in a 100 ms time window around the selected LFP sections were detected. Peaks with maximum amplitudes larger than the LFP mean ± 3SD were defined as sharp event peaks.

#### Modulation of firing rate by cortical sharp events

Sharp LFP event peaks were detected as described above. Firing rate of neurons ± 200 ms around the peaks were calculated in 10 ms bins. Firing rate value averages for bins ± 50 ms around the peaks were compared to the averages for all the other bins (baseline). Neurons were defined as modulated if the average firing rate around the peaks was larger than the mean ± 2SD of the baseline firing rate.

#### Quantification of L5 boutons in the TRN

For investigating L5 bouton distribution in the TRN Thy1-ChR2-EYFP mice (n=3) was used. After perfusion, the brain was removed, and sliced to 50 µm thick coronal slices. To define the boundaries of the TRN, PV staining was carried out (See Method Details/Histology). From each mouse, every fifth slice containing the TRN was used, from which 3 (70 µm x 70 µm x 12 µm) confocal z-stacks were taken, with a 0.12 µm step size (Olympos confocal microscope). The pictures were deconvolved with the Xming software. Stacks were manually sorted into 3 categories: not containing boutons, containing 1-5 boutons (sparse innervation) and containing 5+ boutons (dense innervation).

#### Boxplots

The box of the boxplots show first to the third quartile with a line at the second quartile. The whiskers’ ends label the minimum and maximum values. Average value is marked by x. Values 1.5 times the interquartile range larger than the third quartile or 1.5 times the interquartile range smaller than the first quartile were considered as outliers.

#### Statistics

Statistical comparisons were performed using the Mann-Whitney U test for unpaired sets of data and the Student’s paired sample t-test for paired sets of data. Correlation between two sets of data was measured using Pearson correlation method. Statistical significance was set at 0.05. Results are given as mean ± SEM.

### Mouse Light Neuron Browser

For testing the presence of L5-TRN collaterals at single cell level, we analyzed the single cell reconstruction data of the Mouse Light Neuron Browser database (Chandrashekar, 2017) https://www.janelia.org/open-science/mouselight-neuronbrowser. Cortical neurons with soma position in the L5 and projecting to the thalamus were involved in the analysis. L5 neurons not targeting the thalamus were all found to avoid the TRN and were not involved in the paper. While determining the soma position, anatomical categories were specified to match the nomenclature of our viral tracing experiments (rostral and caudal boundaries in M2 and mPFC were defined at the Br. 1.8 mm level). In each case, we used the raw images to validate the presence of the L5-TRN collaterals in addition to the reconstructed data.

### ADDITIONAL RESOURCES

#### Data and code availability

All data reported in this paper will be shared by the lead contact, László Acsády ’ upon request.

All original code has been deposited at https://github.com and is publicly available as of the date of publication. DOIs are listed in the key resources table.

Any additional information required to reanalyze the data reported in this paper is available from the lead contact upon request

