## Supplementary information for "Region Selective Cortical Control Of The Thalamic Reticular Nucleus"

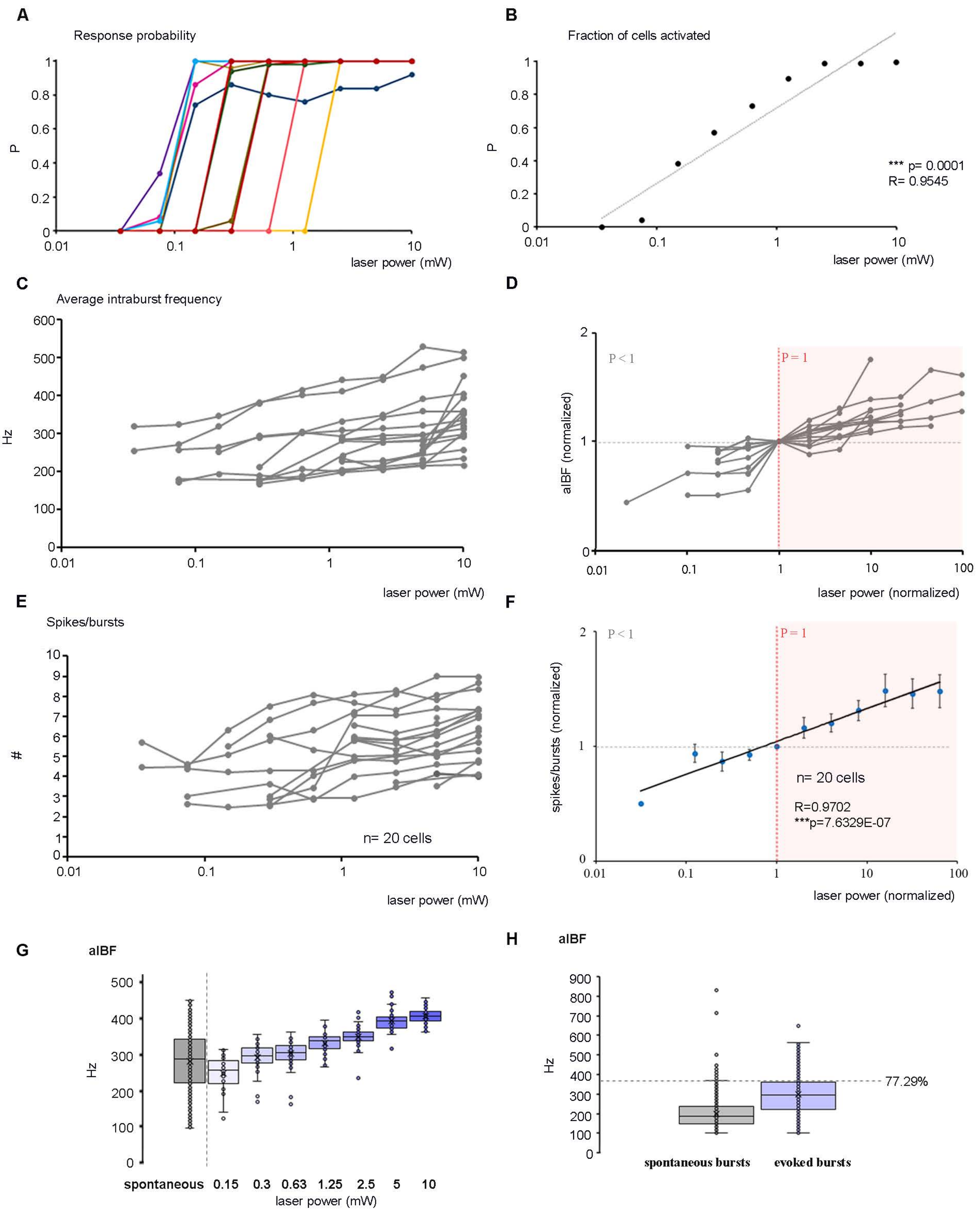


Supplementary figure 1

**Figure S1. All or none response of cortical L5 cells and graded changes in evoked TRN responses at increasing stimulus intensities. Related to Figure 1 and Figure 6.**

(A) Response probabilities of individual cortical L5 cells (n=11) at different laser power values. Laser power is shown on a logarithmic scale.

(B) Population average of response probabilities for cortical L5 cells (n=11) at different laser power values. (Pearson correlation: R=0.9545, *** p=0.0001). Laser power is shown on logarithmic scale.

(C) Average intraburst frequency (aIBF) at different laser power values for individual TRN cells (n=20). Laser power is shown on logarithmic scale.

(D) aIBFs at different laser power values for individual TRN cells (n=20). To compensate the differences between the aIBF ranges and the different number of presynaptic L5 partners recruited at a given laser intensity for each recorded cell, we normalized both the aIBFs and the laser power values to the value observed at the threshold power. Threshold power was the minimal laser power value at which response probability was 1. Laser power is shown on logarithmic scale.

(E) Same as (A) for spikes per burst.

(F) Normalized spike per burst averages for n=20 cells in n=9 mice shown in a log-linear scale. Both the aIBFs and the laser power values are normalized to the value observed at the threshold power. Threshold power was defined as the minimal laser power value at which P=1 response probability was observed. Spike per burst values showed significant log-linear correlation with the laser power.

(G) Box plots for the average intraburst frequencies (aIBFs) of all spontaneous (grey, n=8666 bursts of 17 cells) and evoked (blue, n=4236 bursts of n=17 cells in n=9 mice) TRN bursts. Dashed line indicates the maximum value for the spontaneous bursts. aIBFs of 77.29 % of the optogenetically evoked bursts fall in the range of the values observed during baseline activity.

(H) Box plots for the aIBFs of the spontaneous (grey) and evoked bursts at gradually increasing laser power (blue) for an example cell (baseline: n=649 bursts; 0.15 mW: n=22 bursts; 0.3 mW: n=37 bursts; 0.63 mW: n=49 bursts; 1.25 mW: n=50 bursts; 2.5 mW: n=50 bursts; 5 mW: n=50 bursts).

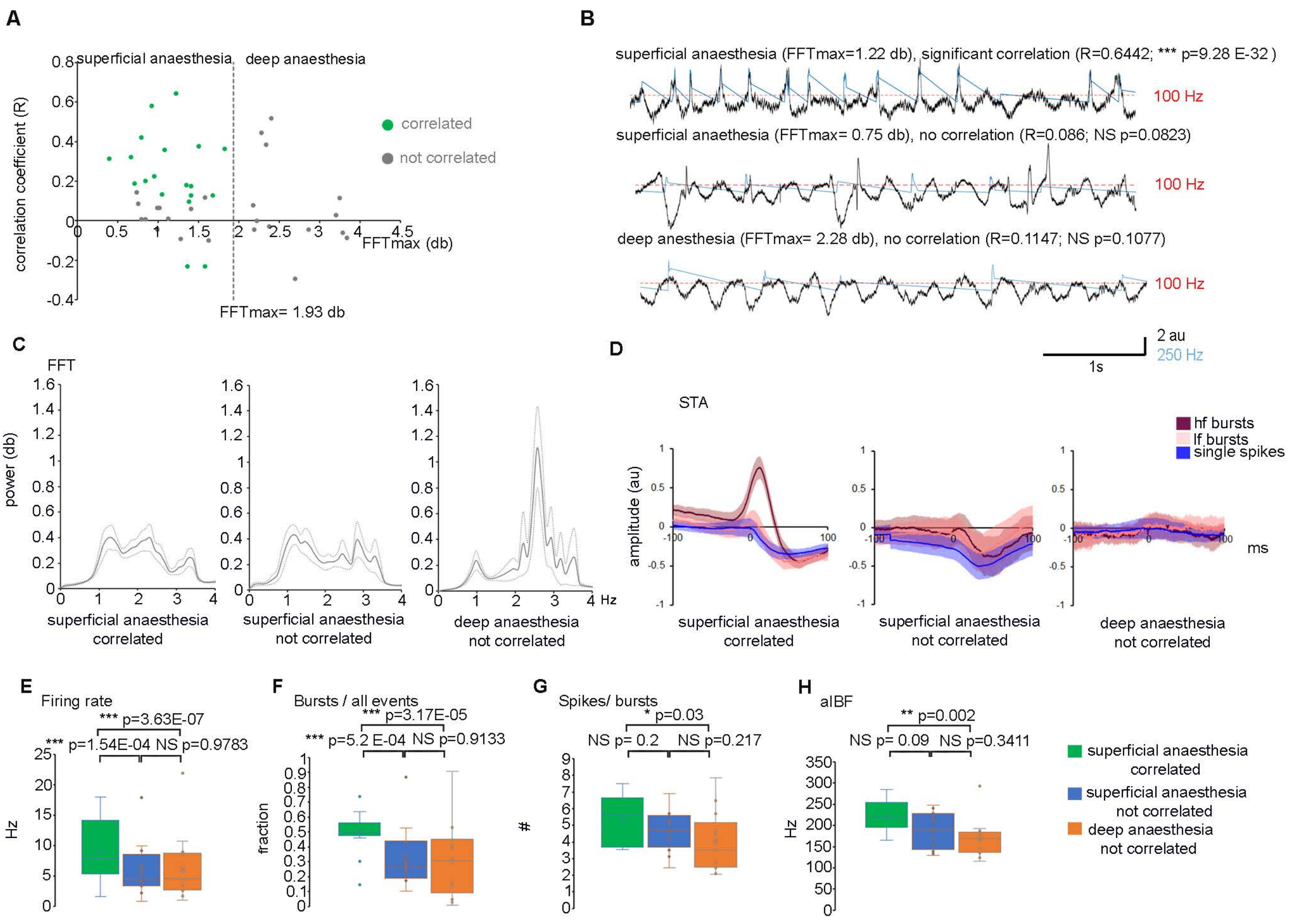


Supplementary figure 2

**Figure S2. Correlation between the spontaneous burst properties and the instantaneous cortical LFP at different depths of anaesthesia. Related to Figure 6.**

(A) Relationship of the aIBF- instantaneous LFP slope correlation coefficient and the peak amplitude of the fast Fourier transform (FFT) of the frontal LFP at the 1-4 Hz range (n=44 cells). Green dots label TRN cells with significant aIBF – instantaneous LFP slope correlation. Grey dots label cells with no significant aIBF- instantaneous LFP slope correlation. Note that when the cortical LFP was dominated by a high amplitude 1-4 Hz component we could not find significant aIBF- instantaneous LFP slope correlation. Superficial – deep anaesthesia boundary (dashed line) was set at average ± 2SD of the FFTmaxs of the 1-4 Hz range using recordings with significant aIBF- instantaneous LFP slope correlation.

(B) Raw frontal LFP traces (black), and instantaneous firing rates (blue) of sample TRN neurons for superficial anaesthesia with significant aIBF- instantaneous LFP slope correlation (upper panel) (FFTmax=1.22 db; R=0.6442; ***p=9.28E-32, n =649 bursts), for superficial anaesthesia with no significant aIBF- instantaneous LFP slope correlation (middle panel) (FFTmax= 0.75 db; R=0.086; NS p=0.0823; n=409 bursts) and for deep anaesthesia with no significant aIBF- instantaneous LFP slope correlation (lower panel) (FFTmax= 2.28 db; R=0.1147; NS p=0.1077; n=198 bursts).
Burst threshold (100 Hz) is labelled by a red dotted line. au: arbitrary unit. Note tight correlation between the cortical LFP and instantaneous firing rate only in the upper panel.

(C) Average±SEM FFT spectrum for recordings with superficial anaesthesia and significant aIBF- instantaneous LFP slope correlation (n=19 cells, left panel), for recordings with superficial anaesthesia and no significant aIBF- instantaneous LFP slope correlation (n=12 cells, middle panel) and for recordings with deep anaesthesia and no significant aIBF- instantaneous LFP slope correlation (n=13 cells, right panel). Note the sharp peak in the delta frequency range on the right panel.

(D) Average LFP values ±SEM (spike triggered averages, STAs) for single spike events (blue), bursts with the lowest 25% of aIBFs (low frequency (lf) burst) (pale red), and bursts with the highest 25% of aIBFs (high frequency (hf) burst) (deep burgundy). Left panel, superficial anaesthesia, significant aIBF- instantaneous LFP slope correlation (n=19 cells); middle panel, superficial anaesthesia, no significant aIBF- instantaneous LFP slope correlation (n=12 cells); right panel, deep anaesthesia, no significant aIBF-instantaneous LFP slope correlation (n=13 cells). HF burst STA peak is only evident in the left panel. au: arbitrary unit.

(E) Firing rate for cells with superficial anaesthesia and significant aIBF- instantaneous LFP slope correlation (9.17±1.17 Hz, n=19 cells) (green), superficial anaesthesia, no significant aIBF- instantaneous LFP slope correlation (6.06±1.32 Hz, n=12 cells) (blue), deep anaesthesia, no significant aIBF- instantaneous LFP slope correlation (6.29±1.53 Hz, n=13 cells) (orange). (Mann-Whitney U Test: superficial, significant aIBF-LFP correlation vs. superficial non-significant aIBF-LFP correlation: *** p=1.54E-04; superficial significant vs. deep: ***p=3.63E-07; superficial non-significant vs. deep: NS p=0.9783)

(F) Same as (E) for burst rate. (superficial anaesthesia, significant aIBF-LFP correlation (n=19 cells): 0.5±0.03; superficial anaesthesia, no significant aIBF-LFP correlation (n=12 cells): 0.34±0.06; deep anaesthesia, no significant aIBF- instantaneous LFP slope correlation (n=13 cells): 0.32±0.07; Mann-Whitney U Test: superficial anaesthesia, significant aIBF-LFP correlation vs. superficial anaesthesia non-significant aIBF-LFP correlation: *** p=5.2 E-04; superficial anaesthesia significant aIBF-LFP correlation vs. deep anaesthesia: *** p=3.17E-05; superficial anaesthesia, no significant aIBF-LFP correlation vs. deep anaesthesia: NS p=0.9133)

(G) Same as (E) for spikes/bursts.

(superficial anaesthesia, significant aIBF-LFP correlation (n=19 cells): 5.36±0.34; superficial anaesthesia, non-significant aIBF-LFP correlation (n=12 cells): 4.67±0.36; deep anaesthesia, not significant aIBF- instantaneous LFP slope correlation (n=13 cells): 3.99±0.5; Mann-Whitney U Test: superficial anaesthesia, significant aIBF – LFP correlation vs. superficial anaesthesia non-significant aIBF- LFP correlation: NS p= 0.2; superficial anaesthesia significant aIBF-LFP correlation vs. deep anaesthesia: * p=0.03; superficial anaesthesia, non-significant aIBF-LFP correlation vs. deep anaesthesia: NS p=0.217)

(H) Same as (E) for aIBF

(superficial anaesthesia, significant aIBF-LFP correlation (n=19 cells): 223.72±8.90; superficial anaesthesia, no significant aIBF-LFP correlation (n=12 cells): 188.85±12.93; deep anaesthesia, no significant aIBF- instantaneous LFP slope correlation (n=13 cells): 170.71±12.44; Mann-Whitney U Test: superficial anaesthesia, significant aIBF – LFP correlation vs. superficial anaesthesia, no significant aIBF-LFP correlation: NS p=0.09; superficial anaesthesia, significant aIBF – LFP correlation vs.

deep anaesthesia: ** p=0.002; superficial anaesthesia, no significant aIBF- LFP correlation vs. deep anaesthesia: NS p=0.3411)

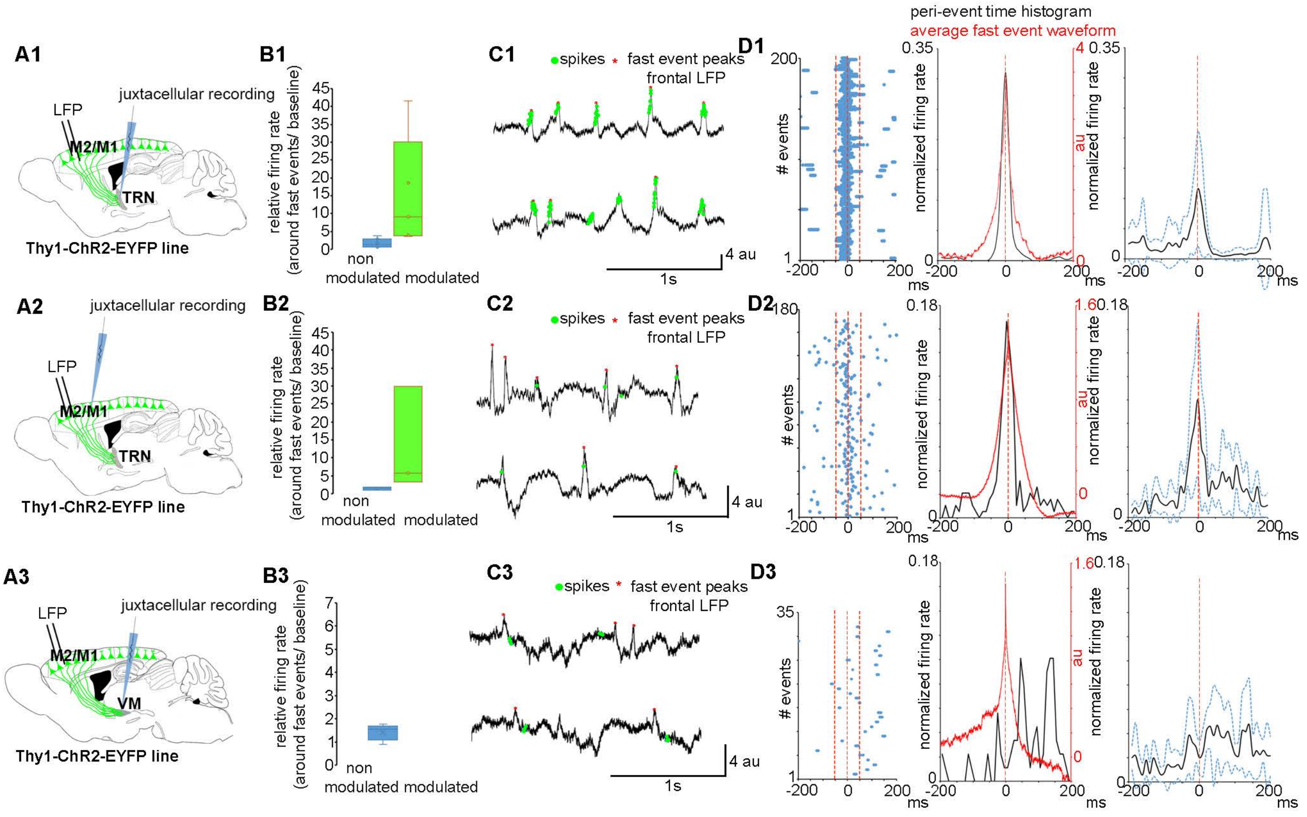


Supplementary figure 3

**Figure S3. Modulation of anterior TRN cell, frontal cortical L5b pyramidal and VM cell firing by fast cortical transients. Related to Figure 6.**

(A1) Schematics of experimental design to record frontal cortical LFP and TRN unit activity.

(A2) Schematics of experimental design to record frontal cortical LFP and L5b unit activity.

(A3) Schematics of experimental design to record frontal cortical LFP and VM unit activity.

(B1) Firing rate of TRN cells around the fast cortical event peaks (±50ms) relative to the baseline activity (50-200 ms before and after the peak). Blue: cells whose firing was not modulated around the peak (n=7 cells) (relative firing rate: 1.78±0.48). Green: cells whose firing was positively modulated around the peak (n=6 cells) (relative firing rate: 15.46±7.09).

(B2) Same as (B1) for L5b cells. (relative firing rate for not modulated cells (n=3): 1.48±0.28; relative firing rate for modulated cells (n=3): 13.04±8.51)

(B3) Same as (B2) for VM cells. (relative firing rate for not modulated cells (n=5): 1.43±0.15)

(D1) Left panel, Spiking activity of an example TRN cell in the ±200ms time window around the peak of the detected fast cortical events; middle panel, peri-event time histogram for an example cell (black). Average fast event waveform (red); right panel, population peri-event time histogram (n=13 cells). Average (black) and ±SEM (blue dotted lines).

(D2) Same as (D1) for L5b cells (n=6 cells)

(D3) Same as (D1) for VM cells (n=5 cells)

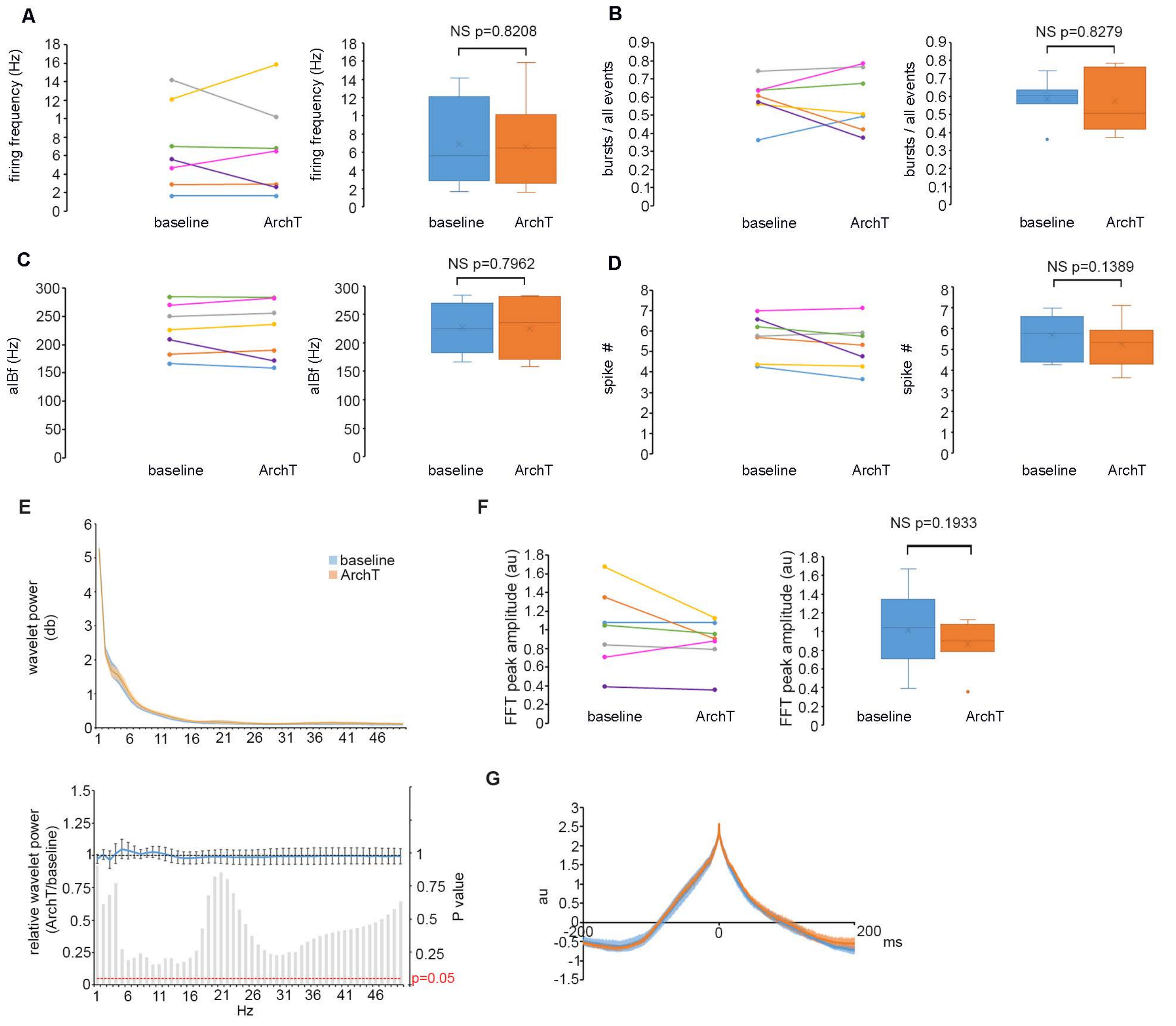


Supplementary figure 4

**Figure S4. Firing properties of the anterior TRN cells and cortical LFP activity did not change during optogenetic perturbation of the L5 inputs in TRN. Related to Figure 7.**

(A) Left panel: Firing frequency of TRN cells at baseline conditions and during optogenetical perturbation of their L5 inputs. Colors of individual cells match the color code on Figure 7. Right panel, box plots of the data in the left panel (6.89±1.76 Hz vs. 6.65±1.9 Hz, Student's paired sample t-test: NS = 0.8208; n= 7 cells). The box of the boxplot shows first to the third quartile with a line at the second quartile. The whiskers' ends label the minimum and maximum values. Average value is marked by x.

(B) Same as (A) for fraction of burst from all firing events. (0.59±0.04 Hz vs. 0.58±0.06 Hz, Student's paired sample t-test: NS p=0.8279; n= 7 cells).

(C) Same as (A) for aIBF. (227.25±16.63 Hz vs. 225.49±19.59 Hz; Student's paired sample t-test: NS p= 0.7962; n= 7 cells).

(D) Same as (A) for spikes per bursts (5.71±0.39 Hz vs. 5.27±0.44 Hz; Student's paired sample t-test: NS p= 0.1389; n= 7 cells)

(E) Upper panel: Average wavelet spectra for recordings of n=7 TRN cells at baseline conditions (blue) and while optogenetically perturbing L5 terminals in the anterior TRN (ArchT)(orange). Average and ± SEM are indicated. Lower panel: average and ± SEM of relative wavelet power during optogenetical perturbation of L5 terminals in the anterior TRN compared to the baseline values (n=7 cells). Box plot below shows significance values for each frequency domain comparing baseline and ArchT values (Student's paired sample t-test, n= 7 cells). 5% significance level is indicated by red dashed line.

(F) Left panel: FFT peak amplitude in the 1-4 Hz range at baseline conditions and during optogenetically inhibiting L5 terminals in the anterior TRN. Colors match color code on Figure 7. Right panel, box plots of the data in the left panel. (1.01±0.16 au vs. 0.87±0.1 au; Student's paired sample t-test: NS p= 0.1933; n= 7 cells) The box of the boxplot shows first to the third quartile with a line at the second quartile. The whiskers' ends label the minimum and maximum values. Average value is marked by x. Values 1.5 times the interquartile range larger than the third quartile or 1.5 times the interquartile range smaller than the first quartile were considered as outliers.

au: arbitrary unit.

(G) Average waveforms of fast cortical events at baseline conditions and during optogenetic perturbation of L5 terminals in the TRN. Average and ± SEM are indicated. (n=7 cells). au: arbitrary unit.

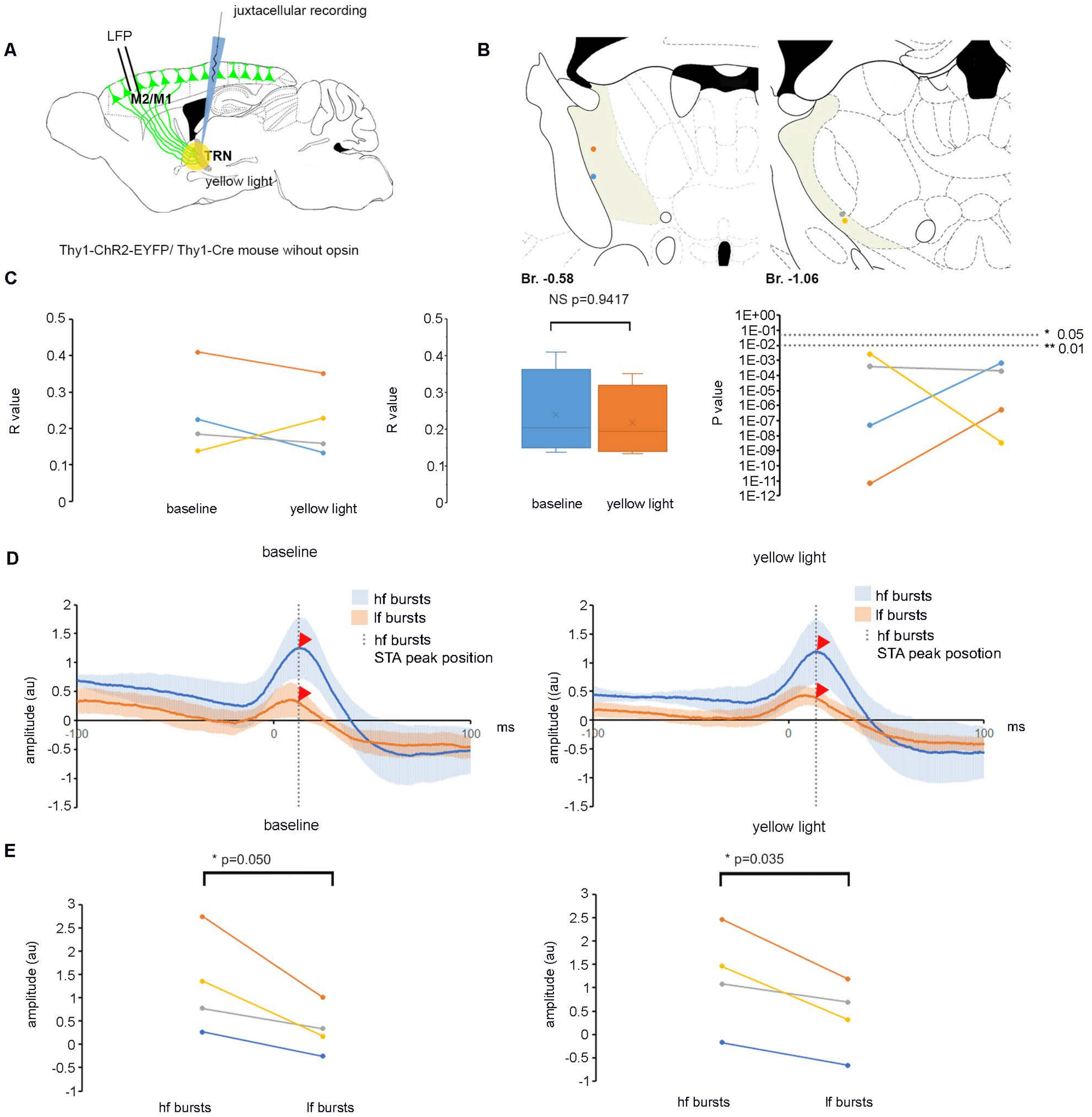


Supplementary figure 5

**Figure S5 Control experiments for the L5-TRN optogenetic perturbation protocol. Related to Figure 7.**

(A) Schematics of experimental design to illuminate anterior TRN with yellow light, and simultaneously record TRN activity and frontal cortical LFP.

(B) Position of juxtacellularly recorded and filled control TRN cells. Colors of the dots match the colors on (C) and (E).

(C) Left panel, Pearson correlation coefficient for the aIBF - LFP slope correlation for n=4 cells at baseline conditions and during optogenetic perturbation protocol with no ChR2 in the L5 fibers (yellow light). Middle panel, box plots of the data in the right panel. (baseline: 0.24±0.06; yellow light: 0.23±0.05); Student's paired sample t-test: NS p=0.9417) The box of the boxplot shows first to the third quartile with a line at the second quartile. The whiskers' ends label the minimum and maximum values. Average value is marked by x. Right panel, P values of Pearson correlation for n=4 TRN cells at baseline vs. during optogenetical perturbation protocol (yellow light). 1% and 5% significance levels are indicated with dashed lines.

(D) Population average of STA traces for n=4 cells. hf bursts (blue); lf bursts (orange). Left panel, baseline; right panel, optogenetic perturbation protocol.
Average and ±SEM are indicated. STA peak for hf bursts is indicated with dashed line. STA amplitudes for hf bursts and lf burst at this value are labelled by red arrowheads. au: arbitrary unit.

(E) STA amplitudes for hf vs. lf bursts at the STA peak for hf bursts at baseline condition (left panel) (hf bursts: 1.28±0.53 au; lf bursts: 0.31±0.26 au; Student's paired sample t-test: * p=0.05) and during optogenetical perturbation protocol (hf bursts: 1.21±0.55 au; lf bursts: 0.38±0.39 au; Student's paired sample t-test: *p=0.035) (yellow light) (right panel). au: arbitrary unit.

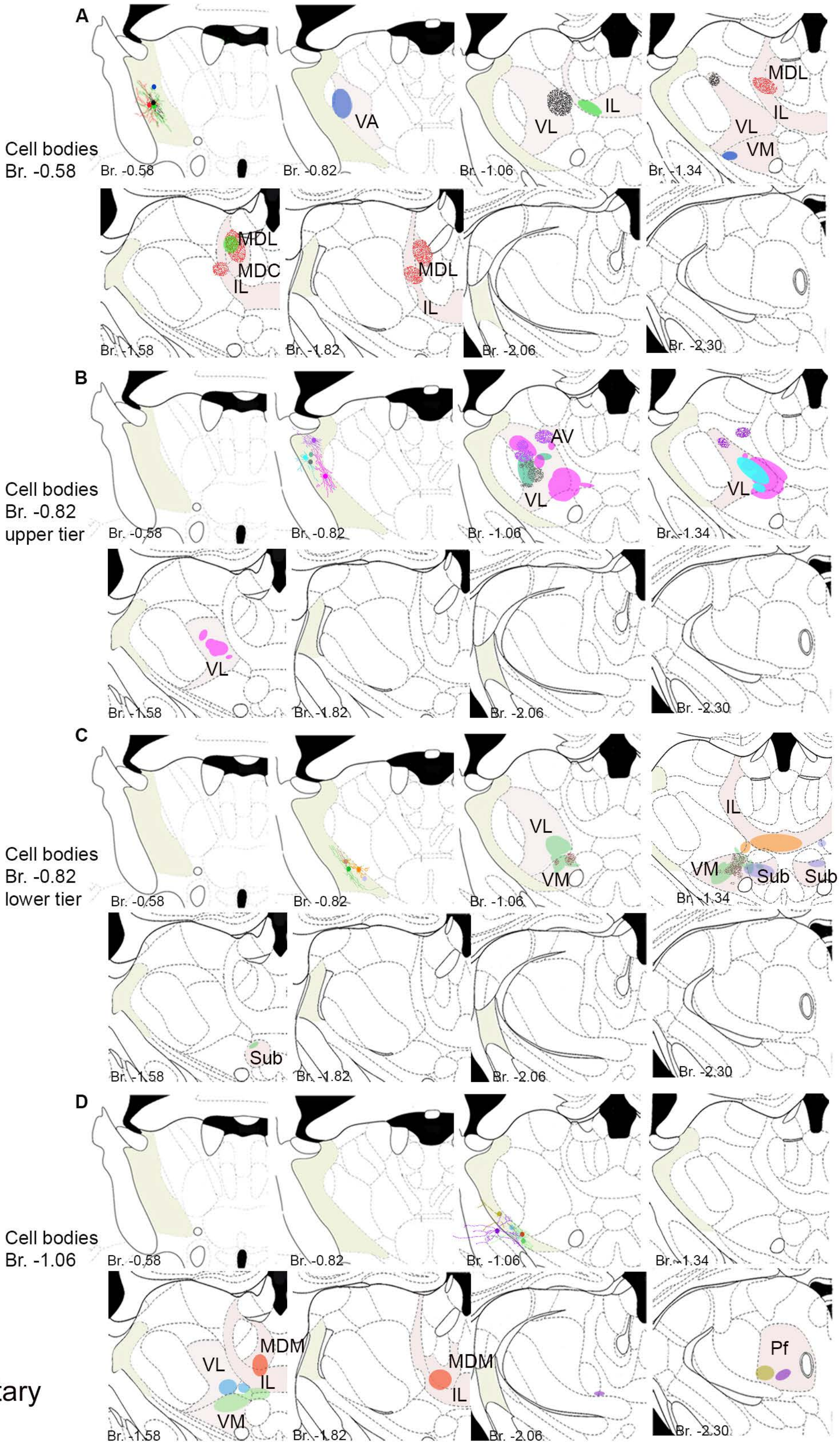


Supplementary figure 6

**Figure S6. Reconstructed dendritic trees, cell body positions and axonal target zone of juxtacellularly recorded and labelled TRN cells optogenetically tagged by their L5 afferents. Related to Figure 8.**

(A) Reconstruction of 4 TRN cells with cell bodies situated at the Br. -0.58 coronal level in eight antero-posterior levels. Dots of different colors within the TRN indicate cell body position. When dendritic trees were filled well enough, it is labelled with the same color as the soma. Axonal target zones in the relay nuclei are indicated with ellipses. Colors of somatic labelling match those of the projection zone. TRN is labelled with pale yellow, target relay nuclei are labelled with pale red. TRN: thalamic reticular nucleus; VA: ventral anterior nucleus; VL: ventral lateral nucleus; IL: intralaminar complex; MDL: mediodorsal nucleus, lateral part; MDC: mediodorsal nucleus, central part; VM: ventral medial nucleus; Br: Bregma

(B) Same as (A) for 5 TRN cells with cell bodies at the Br. -0.82 coronal level, in the upper TRN tier. VL: ventral lateral nucleus; AV: anteroventral nucleus; Br: Bregma

(C) Same as (A) for 4 TRN cells with cell bodies at the Br. -0.82 coronal level in the lower TRN tier. VL: ventral lateral nucleus; VM: ventral medial nucleus; IL: intralaminar complex; Sub: submedius nucleus; Br: Bregma

(D) Same as (A) for 5 TRN cells with cell bodies at the Br. -1.06 coronal level. VL: ventral lateral nucleus; VM: ventral medial nucleus; IL: intralaminar complex; MDM: mediodorsal nucleus, medial part; Pf: parafascicular nucleus Br: Bregma

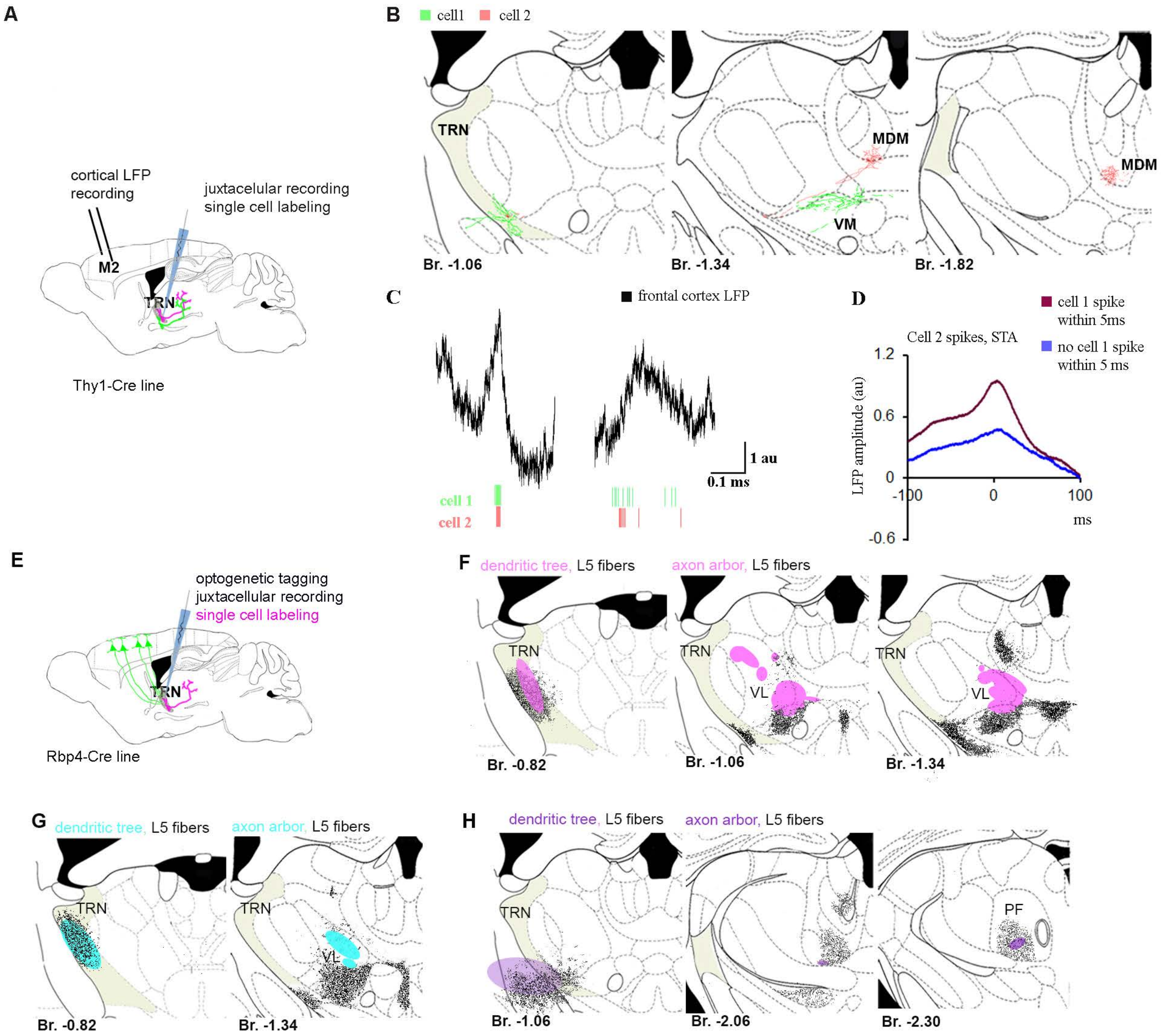


Supplemetary figure 7

**Figure S7. TRN neurons projecting to different relay nuclei synchronize their activity during sharp cortical events and morphological bases for feed forward and lateral inhibition in the L5-TRN pathway. Related to Figure 8.**

(A) Schematics of experimental design for simultaneous recording the baseline activity of two TRN cells and in parallel the LFP activity in the frontal cortex. Both cells were filled with neurobiotin, and their axon arbour was reconstructed.

(B) Left panel: Cell bodies and dendrites of the two TRN cells. For cell one the dendritic tree was only partially labelled. Axon arbour of the two cells in the VM and MDM relay nuclei respectively. TRN, thalamic reticular nucleus; VM, ventral medial nucleus; MDM, mediodorsal nucleus, medial part; Br, Bregma.

(C) Example of firing activity of the two simultaneously recorded TRN cells during a sharp cortical event (left) and during a slower LFP event (right).

(D) Spikes of cell1 were sorted in two groups regarding if they were preceded or followed by a cell2 spike within a 5 ms time window. Spike triggered cortical LFP averages (STAs) were calculated for the two groups. When the two cells fired synchronously (within 5 ms) the STA had higher peak amplitude and larger slope around the spike compared to the STA of spikes when the two cells were less synchronous. au: arbitrary unit.

(E) Schematics of experimental design to label optogenetically tagged and juxtacellularly recorded TRN neurons.

(F) Schematic representation of the dendritic (left panel) and axon (right panel) arbors (magenta) of an example neurobiotin filled TRN neuron optogenetically tagged from L5. Virus labelled L5 fibers are shown in black. Although the dendritic tree highly overlaps with the labelled L5 fibers, TRN axons and L5 collaterals in the relay nuclei do not significantly overlap, suggesting lateral inhibition. TRN: thalamic reticular nucleus; VL: ventral lateral nucleus; Br: Bregma.

(G) Same as (F) for another TRN cell.

(H) Same as (F). In this case, L5 and TRN target zones overlap, implying feed forward inhibition. Dendritic tree of this particular TRN neuron extends to the internal capsule. TRN: thalamic reticular nucleus; PF: parafascicular nucleus; Br: Bregma.

| Frontal cortex | | |
|---|---|---|
| DOI | Soma position | TRN collateral |
| AA0011 | M2 rostral | + |
| AA0114 | M2 rostral | + |
| AA0115 | M1 | + |
| AA0119 | M2 rostral | + |
| AA0121 | M2 rostral | - |
| AA0122 | M2 rostral | + |
| AA0131 | M1 | - |
| AA0132 | M1 | - |
| AA0135 | M2 caudal | - |
| AA0179 | M2 caudal | + |
| AA0181 | M1 | + |
| AA0182 | M2 rostral | - |
| AA0245 | M2 rostral | + |
| AA0250 | M2 rostral | + |
| AA0415 | M2 rostral | + |
| AA0576 | M2 rostral | - |
| AA0583 | M2 caudal | - |
| AA0587 | M1 | + |
| AA0617 | M1 | + |
| AA0644 | M2 caudal | - |
| AA0726 | M2 caudal | + |
| AA0764 | M2 caudal | - |
| AA0772 | M2 rostral | + |
| AA0780 | M2 rostral | - |
| AA0791 | M2 caudal | - |
| AA0792 | M2 caudal | + |
| AA0794 | M2 caudal | + |
| AA0796 | mPFC caudal | + |
| AA0926 | M2 caudal | + |
| AA0927 | M1 | + |
| AA1050 | M1 | + |
| AA1051 | M1 | + |
| Sensory cortices | | |
| AA0001 | S1 | + |
| AA0919 | Aud | - |
| AA0941 | Aud | - |
| AA0944 | S1 | - |
| AA0945 | S1 | - |
| AA0949 | Vis | - |
| AA0956 | S1 | - |
| AA1049 | S1 | - |

Supplementary table 1

**Table S1 Cells involved in the paper from the Mouse Light Neuron Browser database. Related to Figure 1 and Figure 5.**

DOIs and soma position of the reconstructed L5 cortical cells. Presence or absence of TRN collaterals are indicated. The anatomical categories for soma positions were specified to match the nomenclature of our viral tracing experiments.
